# Olig3 acts as a master regulator of cerebellar development

**DOI:** 10.1101/2020.10.10.334615

**Authors:** Elijah D. Lowenstein, Aleksandra Rusanova, Jonas Stelzer, Marc Hernaiz-Llorens, Adrian E. Schroer, Ekaterina Epifanova, Francesca Bladt, Eser Göksu Isik, Shiqi Jia, Victor Tarabykin, Luis R. Hernandez-Miranda

## Abstract

The mature cerebellum controls motor skill precision and participates in other sophisticated brain functions that include learning, cognition and speech. Different types of GABAergic and glutamatergic cerebellar neurons originate in temporal order from two progenitor niches, the ventricular zone and rhombic lip, which express the transcription factors Ptf1a and Atoh1, respectively. However, the molecular machinery required to specify the distinct neuronal types emanating from these progenitor zones is still unclear. Here, we uncover the transcription factor Olig3 as a major determinant in generating the earliest neuronal derivatives emanating from both progenitor zones. In the rhombic lip, Olig3 regulates progenitor cell proliferation. In the ventricular zone, Olig3 safeguards Purkinje cell specification by curtailing the expression of Pax2, a transcription factor that we found to impose an inhibitory interneuron identity. Our work thus defines Olig3 as a key factor in cerebellar development.

## Introduction

The cerebellum develops from the dorsal aspect of rhombomere 1, a region known as the cerebellar anlage that in mice becomes apparent at embryonic (E) day 9.5 (Butts et al., 2014; Chizhikov et al., 2006; Millet et al., 1996; Morales and Hatten, 2006; Wingate and Hatten, 1999). This region contains two distinct germinal zones, the rhombic lip and the ventricular zone, that generate all glutamatergic and GABAergic cerebellar neurons, respectively (Alder et al., 1996; Hallonet et al., 1990; Wingate and Hatten, 1999; Zervas et al., 2004). Development of glutamatergic and GABAergic cerebellar neurons largely depends on the differential expression of two basic helix-loop-helix (bHLH) transcription factors: Atonal homolog 1 transcription factor (Atoh1) and Pancreas specific transcription factor 1a (Ptf1a) (Gazit et al., 2004; Hoshino et al., 2005; Machold and Fishell, 2005; Millen et al., 2014; Wang et al., 2005; Yamada et al., 2014).

In the rhombic lip, Atoh1 directs the generation of three neuronal derivatives: i) deep cerebellar nuclei (DCN) neurons (between E10.5 and E13.5), ii) external granular layer (EGL) cells (between E13.5 and birth), which are the precursors of the internal granular layer cells that develop during early postnatal life, and iii) unipolar brush cells (between E15.5 and the first days of postnatal life) (Ben-Arie et al., 1997; Englund et al., 2006; Fink et al., 2006; Gazit et al., 2004; Machold and Fishell, 2005; Machold et al., 2011; Sekerkova et al., 2004; Yamada et al., 2014). In the ventricular zone, Ptf1a instructs the generation of Purkinje cells (between E11.5-E13.5) and all inhibitory interneurons, including Golgi, Stellate and Basket cells (between E14.5 and birth). Inhibitory interneurons are characterized by the expression of the homeodomain transcription factor Pax2 (Hashimoto and Mikoshiba, 2003; Hoshino et al., 2005; Leto et al., 2006; Maricich and Herrup, 1999). Although the ablation of *Atoh1* and *Ptf1a* severely impairs glutamatergic and GABAergic cerebellar neuron development (Ben-Arie et al., 1997; Hoshino et al., 2005; Jensen et al., 2004; Sellick et al., 2004), less is known about the molecular machinery required for the temporal specification of the different neuronal derivatives emerging from the two cerebellar neurogenic niches.

bHLH transcription factors are master regulators of progenitor cell differentiation during development and are critical players in neuron subtype specification in the nervous system (Atchley and Fitch, 1997; Baker and Brown, 2018; Ben-Arie et al., 2000; Bertrand et al., 2002; Dennis et al., 2019; Dokucu et al., 1996; Imayoshi and Kageyama, 2014; Jones, 2004; Mattar et al., 2008; Ross et al., 2003; Sommer et al., 1996). Among these factors, Oligodendrocyte factor 3 (Olig3) plays a central role in the specification of dorsally emerging neuron types in the hindbrain and spinal cord (Hernandez-Miranda et al., 2017b; Liu et al., 2008; Muller et al., 2005; Storm et al., 2009; Zechner et al., 2007). However, its functions outside these regions have been less studied (Shiraishi et al., 2017; Vue et al., 2007). The combinatorial expression of Olig3 with other bHLH factors, such as Atoh1, Ngn1/2, Ascl1, and Ptf1a, determines the identity of four distinct progenitor cell domains from which defined neuron types emerge in the spinal cord and hindbrain (Hernandez-Miranda et al., 2017a; Liu et al., 2008; Muller et al., 2005; Storm et al., 2009; Zechner et al., 2007). For instance, in the precerebellar system of the hindbrain, Olig3+/Atoh1+ progenitors generate mossy fiber neurons, while Olig3+/Ptf1a+ progenitors generate climbing fiber neurons (Liu et al., 2008; Storm et al., 2009). Although *Olig3* expression has been reported during cerebellar development, its function there has not yet been explored (Liu et al., 2010; Takebayashi et al., 2002).

In this study, we sought to identify bHLH factors that contribute to the development of distinct cerebellar neuron types. We report here that Olig3 is a major regulator of cerebellar development and is crucial for generating the earliest rhombic lip and ventricular zone neuronal derivatives. Our lineage-tracing studies illustrate that the majority of DCN neurons, EGL, granular cells, as well as Purkinje cells emerge from Olig3+ progenitor cells. In contrast, few inhibitory interneurons had a history of *Olig3* expression. Ablation of *Olig3* results in severe cerebellar hypoplasia. In particular, we show that in *Olig3* mutant mice, most DCN neurons as well as half of the EGL cells, granular cells and Purkinje cells are not formed. In contrast, supernumerary inhibitory interneurons develop in *Olig3* mutant animals. Mechanistically, we show that Olig3 cell-autonomously suppresses the development of inhibitory interneurons both in the rhombic lip and the ventricular zone. In the ventricular zone, Olig3 is first expressed in progenitor cells and transiently retained in newborn Purkinje cells to curtail the expression of Pax2, a gene that we found to impose an inhibitory interneuron identity. We also show that Olig3 and its close family member Olig2 specify complementary Purkinje cell populations. Altogether, our data provide new insights into the molecular machinery that secures the correct development of cerebellar neurons.

## Results

### Olig3 is expressed in rhombic lip and ventricular zone progenitor cells during early cerebellar development

To identify candidate bHLH factors that contribute to the generation of early versus late derivatives from the rhombic lip and the ventricular zone, we first analyzed the expression pattern of 108 bHLH transcription factors throughout mouse cerebellar development using publically available data from the Allen Developing Mouse Brain Atlas (https://developingmouse.brain-map.org). We found 50 bHLH genes expressed during cerebellar development, of which 26 were exclusively seen in progenitor niches (rhombic lip, ventricular zone and/or EGL), and the remainder in postmitotic regions (Figure 1A and Table S1). In particular, 9/26 genes displayed differential spatial-temporal expression patterns in the rhombic lip (Atoh1 and Olig3 between E11.5-E13.5), EGL (Atoh1 and Neurod1 between E13.5-birth) and ventricular zone (Ptf1a, Ascl1, Olig3 and Olig2 between E11.5-E13.5; Ptf1a, Neurod6, Neurog1 and Neurog2 between E13.5-E18.5). The remaining (17/26) factors appeared to belong to either a common set of transcription factors expressed in all progenitor niches or they maintained their expression in a particular niche throughout cerebellar development (Table S1). Of particular interest was the expression pattern of Olig3, which has not been previously reported to have a function in cerebellar development but is essential for the specification of numerous hindbrain and spinal cord neurons (Hernandez-Miranda et al., 2017a; Liu et al., 2008; Muller et al., 2005; Storm et al., 2009; Zechner et al., 2007).

**Figure 1.**
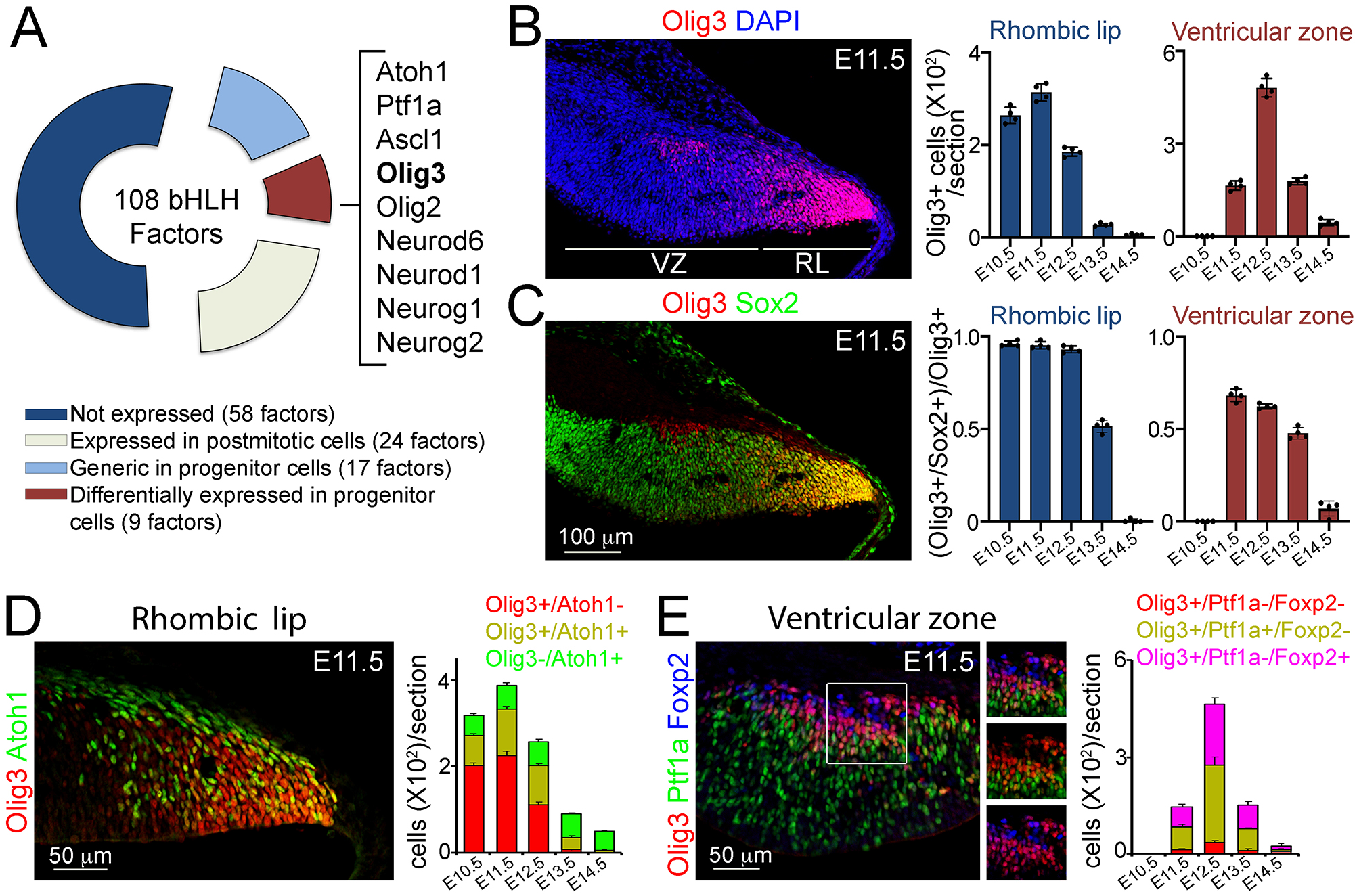
Olig3 marks rhombic lip and ventricular zone progenitor cells during early cerebellar development. (**A**) Doughnut chart illustrating the expression of 108 bHLH transcription factors during cerebellar development. For details on individual gene names within each category please see Table S1. (**B, C**) Left, a sagittal section of the cerebellum stained with antibodies against Olig3 (red) and counterstained with DAPI (blue in B) and with antibodies against Sox2 (green in C) at E11.5. Right, quantification of Olig3+ cells (upper graphs) and the proportion of Olig3+ cells co-expressing Sox2 (lower graphs) in the rhombic lip (RL) and ventricular zone (VZ) at different embryonic days. See also Figure S1A. Dots in the graphs represent the mean of individual analyzed animals. (**D**) Left, immunofluorescence characterization of rhombic lip progenitor cells stained against Olig3 (red) and Atoh1 (green) at E11.5. Right, quantification of the proportion of Olig3+ rhombic lip cells co-expressing Atoh1 at different embryonic days. (**E**) Left, immunofluorescence characterization of ventricular zone progenitor cells stained against Olig3 (red), Ptf1a (green) and the Purkinje cell marker Foxp2 (blue) at E11.5. Right, quantification of the proportion of Olig3+ ventricular zone cells co-expressing Ptf1a or Foxp2. The mean and SD are plotted in all graphs. Photomicrographs were acquired using the automatic tile scan modus (10% overlap between tiles) of the Zeiss LSM700 confocal microscope.

Next, we characterized the expression of *Olig3* during cerebellar development by immunofluorescence. In the rhombic lip, Olig3+ cells were abundant from E10.5 to E12.5, but their numbers declined by E13.5 and were rare by E14.5 (Figures 1B and S1A). Almost all (>98%) Olig3+ rhombic lip cells co-expressed the progenitor marker Sox2 between E10.5 and E12.5, but this co-localization, as well as the total number of Olig3+ cells, declined by E13.5 (Figure 1C). In addition, most (>95%) proliferative BrdU+ cells in the rhombic lip co-expressed Olig3 (Figure S1B). Lastly, about 30% of Olig3+ cells in the rhombic lip expressed Atoh1 (Olig3+/Atoh1+ cells; Figure 1D). Thus, in the rhombic lip, Olig3+ cells are progenitors, and a third of them co-express Atoh1.

In the ventricular zone, Olig3+ cells were first seen at E11.5. Their numbers peaked by E12.5 and became rare by E14.5 (Figures 1B and S1A). Most Olig3+ cells (59%) in the ventricular zone co-expressed the progenitor marker Sox2 at E11.5 and E12.5, but this co-localization, as well as the total number of Olig3+ cells, declined by E13.5 (Figure 1C). Furthermore, about one-third of the BrdU+ cells in the ventricular zone co-expressed Olig3 (Figure S1C). Lastly, 52% of the Olig3+ ventricular zone cells co-expressed Ptf1a, while 41% co-expressed the postmitotic Purkinje cell marker Foxp2 (Figure 1E). This indicates that whereas most Olig3+ cells in the ventricular zone are progenitors (Olig3+/Ptf1a+/Foxp2-), Olig3 is transiently retained in early-born Purkinje cells (Olig3+/Ptf1a-/Foxp2+). We conclude that Olig3 is expressed in rhombic lip and ventricular zone progenitor cells during the generation of their earliest neuronal derivatives.

### Early derivatives from the rhombic lip and ventricular zone arise from Olig3+ progenitor cells

To obtain a complete picture of the distinct cerebellar neuron types arising from Olig3+ progenitor cells, we first carried out a long-term lineage tracing experiment. This experiment used a tamoxifen-inducible cre recombinase driven by *Olig3 (Olig3^creERT2/+^*) and the fluorescent reporter *Ai14* that expresses a cytoplasmic Tomato fluorescent protein upon cre-mediated recombination (see the genetic strategy in Figure S2A). Specifically, we induced tamoxifen recombination in *Olig3^creERT2/+^;Ai14^+/-^* mice at E10.5 and analyzed their brains by light-sheet microscopy at E19 (see Video S1). Three-dimensional reconstructions of recombined brains showed that Tomato+ cells were broadly distributed across the entire cerebellum of *Olig3^creERT2/+^; Ai14^+/-^* mice (Figures 2A and S2A). In particular, we observed Tomato+ cells in the EGL, Purkinje cell layer, and dense groups of Tomato+ cells encompassing the three nuclei formed by DCN neurons: the nucleus dentatus, interpositus, and fastigii (Figures 2A-C). Closer inspection revealed that Tomato+ cells co-expressed Tbr1 (a marker of Fastigii DCN cells) and Brn2 (a marker of Interpositus and Dentatus DCN cells) (Figure 2D). Thus, the distribution of cerebellar neurons with a history of *Olig3* expression is compatible with the notion that Olig3+ progenitor cells generate the earliest set of glutamatergic (DCN, EGL) and GABAergic (Purkinje cells) cerebellar derivatives.

**Figure 2.**
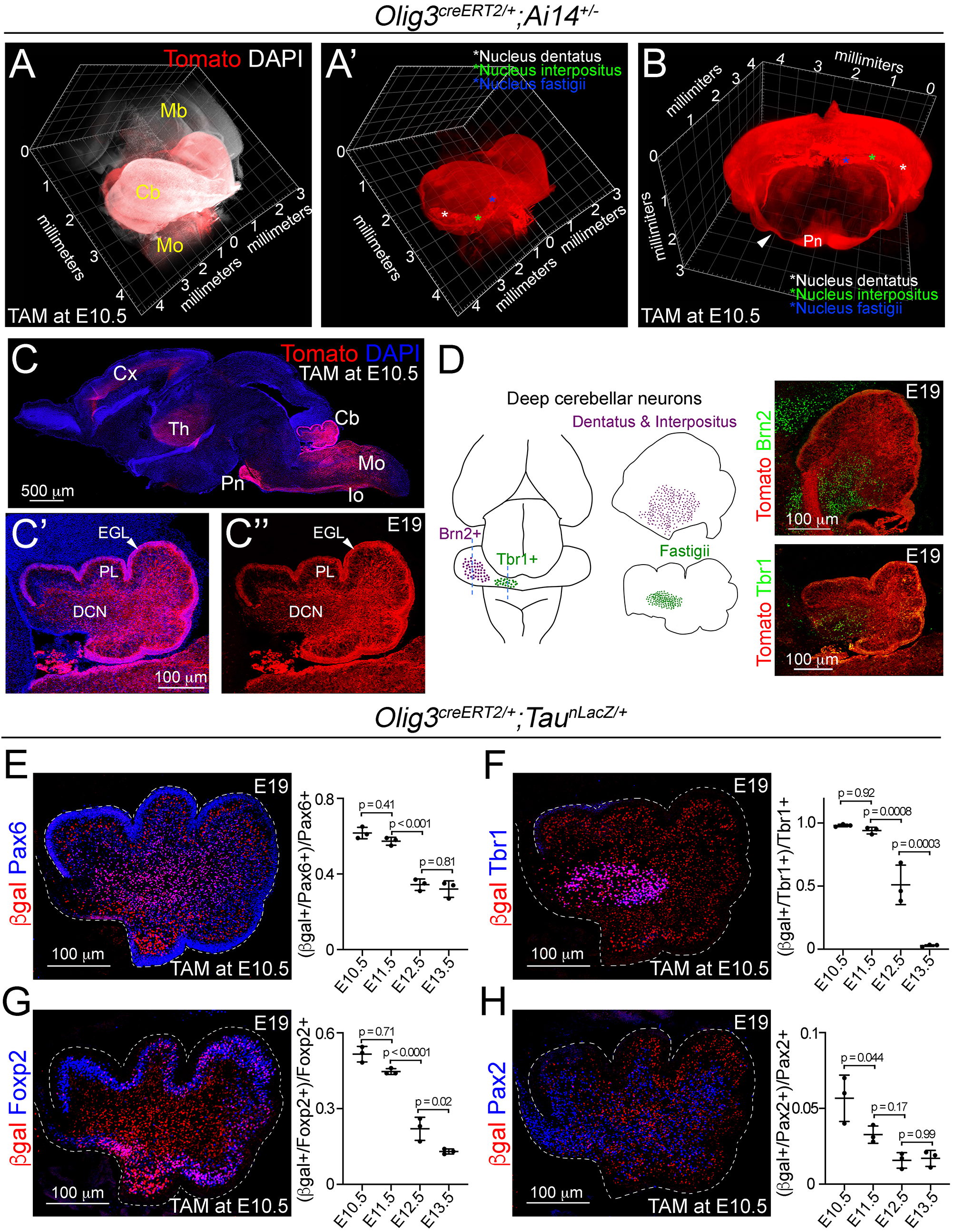
Lineage-tracing of cerebellar neurons arising from Olig3+ progenitor cells. (**A-D**) Analysis of Tomato+ (red) cells in *Olig3^creERT2/+^;Ai14^+/-^* mice that were recombined with tamoxifen (TAM) at E10.5 and imaged at E19. The *Ai14* allele encodes for a cytoplasmic red fluorescent protein that labels both cell bodies and axons. See Figure S2A and Video S1 for a complete reconstruction of a recombined *Olig3^creERT2/+^;Ai14^+/-^* brain. (**A, A’**) A sagittal three-dimensional reconstruction of the cerebellum. Tomato+ cells were broadly distributed across the entire cerebellum and densely packed in the three DCN nuclei (asterisks in A’). (**B**) A coronal three-dimensional reconstruction of the cerebellum. DCN nuclei are marked with asterisks. The pontine nuclei (Pn) and their axons (arrowhead), which develop from Olig3+ progenitor cells in the medulla oblongata, are labeled with Tomato. (**C**) A sagittal section stained against Tomato and DAPI (blue). Other known Olig3 derivatives, such as the thalamus (Th) including its projections to the cortex (Cx), pontine nuclei (Pn), inferior olive (Io) and many neurons in the medulla oblongata (Mo) are also marked with Tomato. A magnification of the cerebellum is displayed with (C’) or without (C”) DAPI. The external granular cell layer (EGL), Purkinje cell layer (PL) and DCN neurons are labeled with Tomato. (**D**) Left, schematic display of DCN nuclei that are positive for Brn2 (dentatus and interpositus) and Tbr1 (fastigii). Right, sagittal cerebellar sections stained against Tomato and Brn2 or Tbr1 (green). (**E-H**) Analysis of cerebellar neurons with a history of Olig3 (βgal+) expression. *Olig3^creERT2/+^;Tau^nLacZ/+^* mice were recombined with tamoxifen (TAM) at different embryonic stages and analyzed at E19. See Figure S2B for a description of the experiment. Sagittal cerebellar sections from these mice were stained against βgal (red) and markers for the EGL and granular cells (Pax6, blue in E), DCN neurons (Tbr1, blue in F), Purkinje cells (Foxp2, blue in G) and inhibitory interneurons (Pax2, blue in H). Double-positive (βgal+/marker+) cells were quantified at E19. The mean and SD are plotted in all graphs, and the dots represent the mean of individual animals. Significance was obtained using one-way ANOVA followed by *post hoc* Tukey’s test, see Table S2. Photomicrographs were acquired using the automatic tile scan modus (10% overlap between tiles) of the Zeiss LSM700 confocal microscope.

We performed a second long-term lineage tracing experiment to better define the temporal contribution of Olig3+ progenitor cells to specific cerebellar neuron types. In particular, we used *Olig3^creERT2^* and the reporter *Tau^nLacZ^,* which selectively expresses a nuclear β-galactosidase (βgal) protein upon cre-mediated recombination in postmitotic (Tau+) neurons. We induced tamoxifen recombination in *Olig3^creERT2/+^;Tau^nLacZ/+^* mice at distinct embryonic stages from E10.5 to E13.5 and analyzed the cerebella of recombined mice at E19 (Figure S2B-C). As expected, no βgal+ cell was observed in the EGL, as this layer contains granular cell progenitors that do not express Tau (arrowheads in Figure S2C). Both EGL and postmitotic granular cells express the transcription factor Pax6 (Fink et al., 2006; Yeung et al., 2016). We observed that the majority (62%) of Pax6+ postmitotic cells, outside the EGL, co-expressed βgal when recombination was induced at E10.5 or E11.5, but the proportion of Pax6+/βgal+ cells dropped when recombination was induced at later stages (Figure 2E). Furthermore, we found that most Tbr1+ (>99%) DCN neurons, Foxp2+ (51%) Purkinje cells, and Brn2+ (43%) DCN neurons co-expressed βgal when recombination was induced at E10.5 or E11.5, but the proportion of double-positive cells declined when recombination was induced at later stages (Figures 2F, 2G, and S2D). In contrast, few Pax2+ (6%) cells co-expressed βgal when recombination was induced at E10.5, and the number of double positive cells was minimal when recombination was induced at later stages (Figure 2H). Unipolar brush cells (Tbr2+) were also observed to co-express βgal in recombined animals (Figure S2E). One should note that these cells derive from late rhombic lip progenitor cells at a time point (E15.5-birth) when *Olig3* is no longer expressed, which indicates that these progenitor cells, like those in the EGL, had a history of *Olig3* expression. We conclude that Olig3+ progenitor cells substantially contribute to the generation of the earliest derivatives of the rhombic lip (DCN and EGL cells) and the ventricular zone (Purkinje cells).

### Cerebellar hypoplasia and loss of early-born cerebellar neurons in *Olig3* mutant mice

We next analyzed the consequences of *Olig3* ablation on cerebellar development. At birth (P0), the cerebella of *Olig3* null *(Olig3^-/-^*) mutant mice were drastically reduced in volume when compared to control *(Olig3^+/-^*) littermates (Figures 3A, 3B). The strongest reduction in volume was observed in the medial portion of the cerebella of *Olig3* mutant mice (Fig. S3A). Closer inspection of *Olig3* mutant cerebella revealed that they had less folia than control littermates (Figure 3B). Thus, ablation of *Olig3* results in severe cerebellar hypoplasia. Furthermore, the number of Tbr1+, Brn2+, Pax6+ and Foxp2+ neurons was greatly reduced in *Olig3* mutant mice (Figures 3C-E and S3B). In contrast, the number of Pax2+ inhibitory interneurons increased in mutant mice (Figure 3F). Late born derivatives from the rhombic lip, such as Tbr2+ unipolar brush cells, were correctly specified in *Olig3* mutant mice (Figure S3C). Together these data show that Olig3 is critically involved in cerebellar development and the generation of DCN neurons, EGL cells (including their granular cell derivatives) and Purkinje cells.

**Figure 3.**
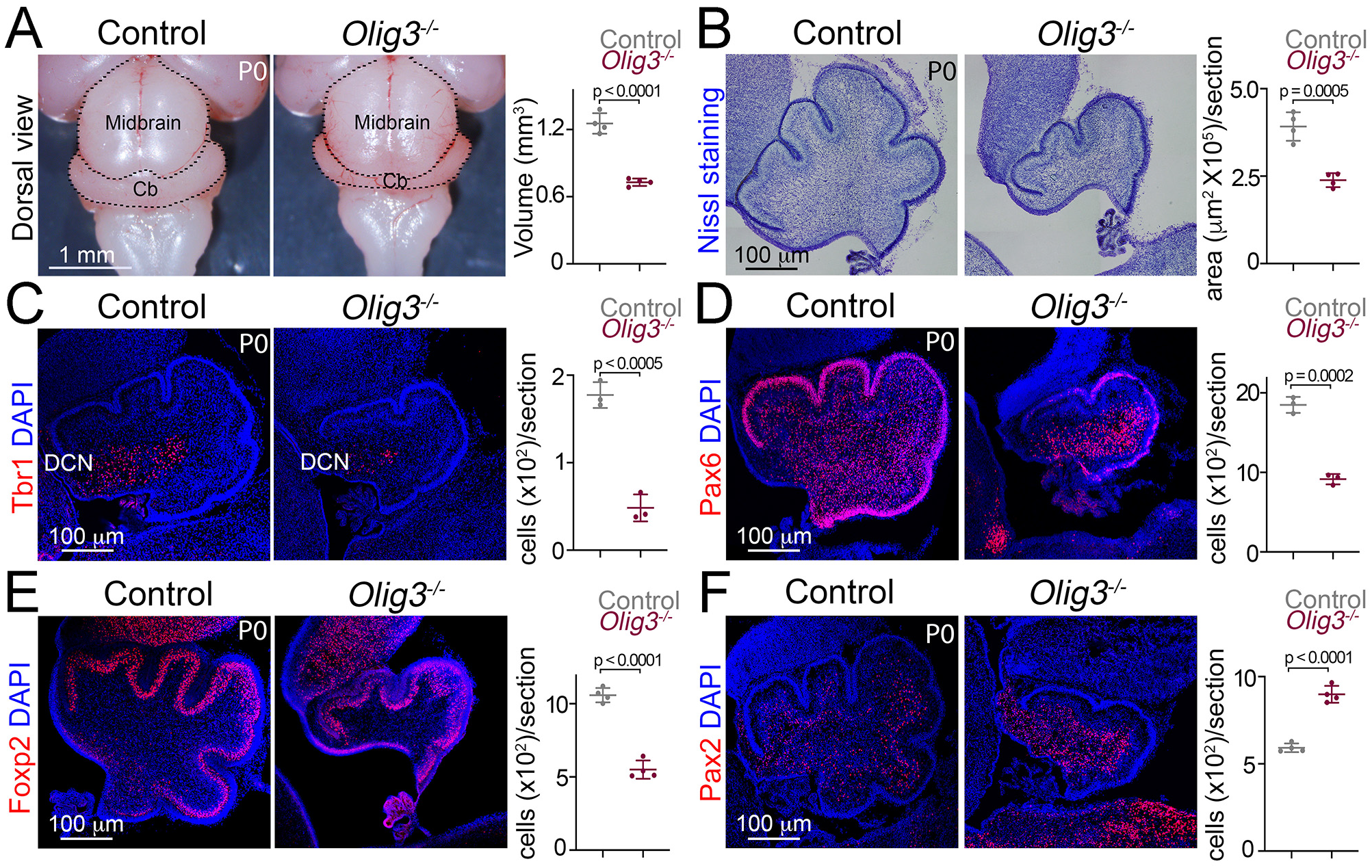
Severe cerebellar hypoplasia and neuronal loss in *Olig3* mutant mice. (**A**) Left, dorsal views of control *(Olig3^+/-^*) and *Olig3* mutant *(Olig3^-/-^*) cerebella at birth (P0). Right, quantification of cerebellar volume in newborn control and *Olig3^-/-^* mice. (**B**) Left, sagittal sections of newborn control and *Olig3^-/-^* cerebella stained with Nissl. Right, quantification of cerebellar area in newborn control and *Olig3^-/-^* mice. (**C-F**) Immunofluorescence characterization and quantification of Tbr1+ DCN neurons (C, in red), Pax6+ EGL and granular cells (D, in red), Foxp2+ Purkinje cells (E, in red) and Pax2+ inhibitory interneurons (F, in red) in newborn control and *Olig3^-/-^* mice. All cerebellar sagittal sections were counterstained with DAPI (blue). The mean and SD are plotted in all graphs, and the dots represent the mean of individual animals. Two-tailed t-tests were performed to determine statistical significance. See Table S2. Photographs in A and B were acquired with a conventional bright-field microscope and photomicrographs in C-F were acquired using the automatic tile scan modus (10% overlap between tiles) of the Zeiss LSM700 confocal microscope.

To define the developmental onset of the cerebellar deficiencies seen in *Olig3* mutant mice, we carried out a short-term lineage tracing experiment using a knock-in mouse strain that expresses GFP from the *Olig3* locus (*Olig3^GFP^*) (Muller et al., 2005). We compared control (*Olig3^GFP/+^*) and *Olig3* mutant (*Olig3^GFP/GFP^*) mice at E13.5. In control mice at this stage, DCN neurons have already migrated away from the rhombic lip and accumulated in the nuclear transitory zone, EGL cells have formed their characteristic subpial layer, and Purkinje cells have completed their specification (Figure S4). Compared to control littermates, E13.5 *Olig3* mutant mice showed a severe reduction in the number of DCN and EGL cells but no significant reduction in the number of Purkinje cells (Figure S4). Therefore, while the deficits observed in DCN neurons and EGL cells arise early during cerebellar development in *Olig3* mutant mice, the reduction in Purkinje cell numbers occurs at a later developmental stage than E13.5 (see below).

We next analyzed whether Atoh1 and/or Ptf1a progenitor cell numbers changed in *Olig3* mutant mice at early embryonic stages. Analysis of *Olig3* mutant mice at E11.5 and E12.5 revealed reduced numbers of Atoh1+ cells in the rhombic lip, but no change in the number of Ptf1a+ cells in the ventricular zone (Figures S5A and S5B). We then asked whether the ablation of *Olig3* might affect the proliferation of rhombic lip and ventricular zone progenitor cells and/or their viability. In the rhombic lip, the number of proliferative (BrdU+) cells was reduced in *Olig3* mutant animals. However, no change in the number of Tunel+ apoptotic bodies (puncta) were seen at any of the analyzed embryonic stages (Figures S5C and S5D). In the ventricular zone of *Olig3* mutant animals neither the number of BrdU+ or Tunel+ apoptotic bodies changed (Figures S5C and S5D). Thus, the mutation of *Olig3* impairs progenitor proliferation in the rhombic lip but not in the ventricular zone, illustrating that Olig3’s function in the ventricular zone differs from that in the rhombic lip.

### Ablation of *Olig3* misspecifies Purkinje cells that transform into inhibitory interneurons

We next compared the development of Foxp2+ Purkinje cells and Pax2+ inhibitory interneurons in *Olig3* mutant mice. In wildtype and heterozygous *Olig3^GFP/+^* mice, Pax2+ cells first appeared at E13.5 in a rostral domain of the ventricular zone that lacked expression of Olig3, and by E14.5 occupied most of the ventricular zone (Figure 4A). The spread of Pax2+ cells from rostral to caudal coincided with the receding of Olig3+ cells (schematically displayed in Figure 4B). In sharp contrast to wildtype and heterozygous *Olig3^GFP/+^* mice, we found supernumerary Pax2+ cells in *Olig3^GFP/GFP^* mutant mice from E13.5 to P0 (Figures 4C and S6A, quantified in Figure 4D). Many of these co-expressed GFP in the ventricular zone and rhombic lip at E13.5 and E14.5 (see magnifications in Figures 4C and S6A). Thus, the ablation of *Olig3* derepresses Pax2 in the ventricular zone and the rhombic lip.

**Figure 4.**
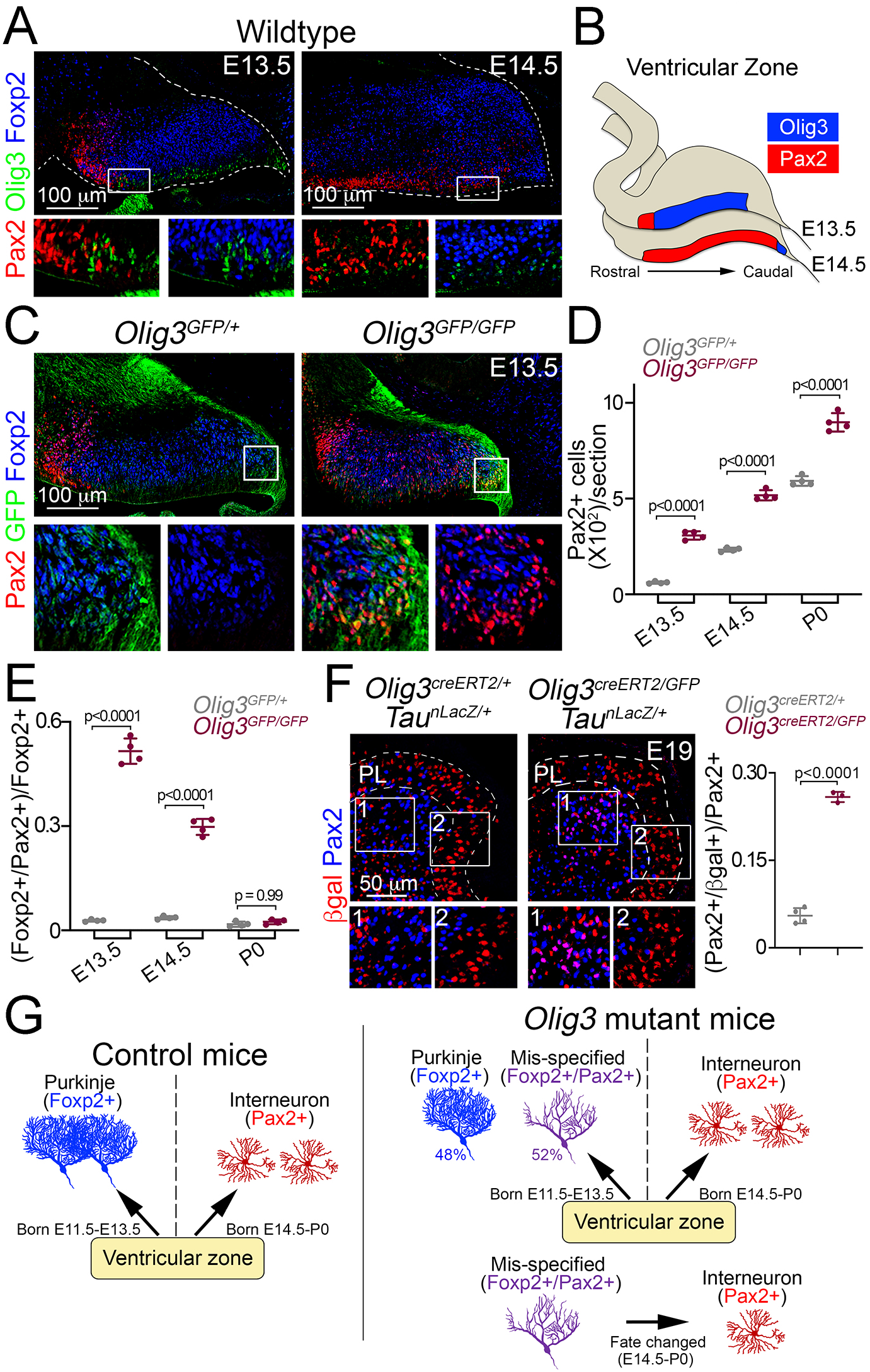
Ablation of *Olig3* misspecifies Purkinje cells which become inhibitory interneurons. (**A**) Immunofluorescence characterization of Foxp2+ (blue) Purkinje cells, Pax2+ (red) inhibitory interneurons and Olig3+ (green) progenitor cells in wildtype mice at the indicated stages. Boxed areas are magnified underneath the main photographs. (**B**) Schema illustrating the development of Pax2+ inhibitory interneurons. At E13.5, inhibitory interneurons develop in a rostral domain of the ventricular zone that lacks Olig3 expression. At E14.5, Pax2+ cells span most of the ventricular zone as Olig3 expression becomes extinguished. (**C**) Immunofluorescence characterization of Foxp2+ (blue) Purkinje cells and Pax2+ (red) inhibitory interneurons in E13.5 control (*Olig3^GFP/+^*) and *Olig3* mutant (*Olig3^GFP/GFP^*) mice. All cerebellar sagittal sections were stained against GFP (green). Boxed areas are magnified underneath the main photographs. Note that GFP+ and Foxp2+ cells ectopically express Pax2 in *Olig3^GFP/GFP^* mice. (**D**) Quantification of Pax2+ cells in *Olig3^GFP/+^* and *Olig3^GFP/GFP^* mice at the indicated stages. (**E**) Quantification of the proportion of Foxp2+ Purkinje cells co-expressing Pax2+ in *Olig3^GFP/+^* and *Olig3^GFP/GFP^* mice at the indicated stages. (**F**) Immunofluorescence characterization and quantification of the proportion of Pax2+ (blue) inhibitory interneurons co-expressing βgal (red) in E19 control (*Olig3^creERT2/+^;Tau^nLacZ/+^*) and *Olig3* mutant (*Olig3^creERT2/GFP^;Tau^nLacZ/+^*) mice that were recombined at E10.5. (**G**) Schema illustrating the above findings. In control mice, the ventricular zone generates two sets of GABAergic neurons: Foxp2+ Purkinje cells (E11.5-E13.5) and Pax2+ inhibitory interneurons (E14.5-P0). In *Olig3* mutant mice, about half of the Foxp2+ cells are misspecified and co-expressed Pax2. These cells subsequently change their fate and transform into inhibitory interneurons. The mean and SD are plotted in all graphs, and the dots represent the mean of individual animals. Significance was determined using a one-way ANOVA followed by *post hoc* Tukey’s (in D and E) or two-tailed t-test (in F) analyses, see Table S2. Photomicrographs were acquired using the automatic tile scan modus (10% overlap between tiles) of the Zeiss spinning disk confocal microscope (in A and *C*) and the Zeiss LSM700 confocal microscope (in F).

Surprisingly at E13.5, most Foxp2+ cells (52%) co-expressed Pax2 in *Olig3* mutant mice (Figure 4C, quantified in Figure 4E), vice versa, roughly 90% of the Pax2+ cells co-expressed Foxp2 (Figure S6B). Interestingly, the proportion of misspecified Foxp2+/Pax2+ (or Pax2+/Foxp2+) cells declined by E14.5 and became rare by P0 (quantified in Figures 4E and S6B). The decrease of mis-specified (Foxp2+/Pax2+) cells coincided with the increase of inhibitory (Foxp2-/Pax2+) interneurons seen in *Olig3* mutant animals (compare Figures 4D and 4E). We thus hypothesized that misspecified (Foxp2+/Pax2+) cells in *Olig3* mutant animals might undergo a fate shift and adopt an inhibitory (Foxp2-/Pax2+) interneuron identity. To assess this hypothesis, we carried out a long-term lineage tracing experiment using *Olig3^creERT2^* and the *Tau^nLacZ^* reporter in an *Olig3* mutant background (*Olig3^creERT2/GFP^;Tau^nLacZ/+^* mice) and analyzed βgal expression in Pax2+ inhibitory interneurons. Tamoxifen recombination in *Olig3^creERT2/GFP^;Tau^nLacZ/+^* mice was induced at E10.5. We found an increase in the proportion of Pax2+/ βgal+ cells in *Olig3^creERT2/GFP^;Tau^nLacZ/+^* mice when compared to *Olig3^creERT2/+^;Tau^nLacZ/+^* control littermates (Figure 4F). Notably, ectopic Pax2+/ βgal+ cells in *Olig3^creERT2/GFP^;Tau^nLacZ/+^* mice adopt an inhibitory interneuron identity (Pax2 expression), and also intermingle with Pax2+/ βgal-cells underneath the Purkinje cell layer at E19 (compare inserts 1 and 2 in Figure 4F). Taken together, we conclude that ablation of *Olig3* in the ventricular zone results in the misspecification of Purkinje cells. These cells later change their fate and adopt an inhibitory interneuron identity (schematically display in Figure 4G).

### Pax2 imposes an inhibitory interneuron indentity that is cell-autonomously suppressed by Olig3

To experimentally assess whether Pax2 cell-autonomously suppresses *Foxp2* expression to impose an inhibitory interneuron fate, we electroporated *in utero* a *Pax2-IRES-GFP* expressing vector in the ventricular zone of wildtype animals at E12.5. This is a timepoint during which Foxp2+ cells are abundant and Pax2+ cells are absent (see Figure 5A for a schematic display of the experimental conditions). Electroporated *pCAG-Pax2-IRES-GFP (Pax2* overexpressing) and *pCAG-GFP* + *Empty-IRES-GFP* (control) embryos were analyzed at E14.5. In *pCAG-Pax2-ires-GFP* electroporated mice, no GFP+/Pax2+ cell co-expressed Foxp2, whereas in *pCAG-GFP* + *Empty-IRES-GFP* electroporated mice about 72% of the GFP+ cells were also Foxp2+ (Figure 5B). We conclude that Pax2 is an efficient selector gene that imposes an inhibitory interneuron fate in derivatives of the ventricular zone by suppressing *Foxp2.*

**Figure 5.**
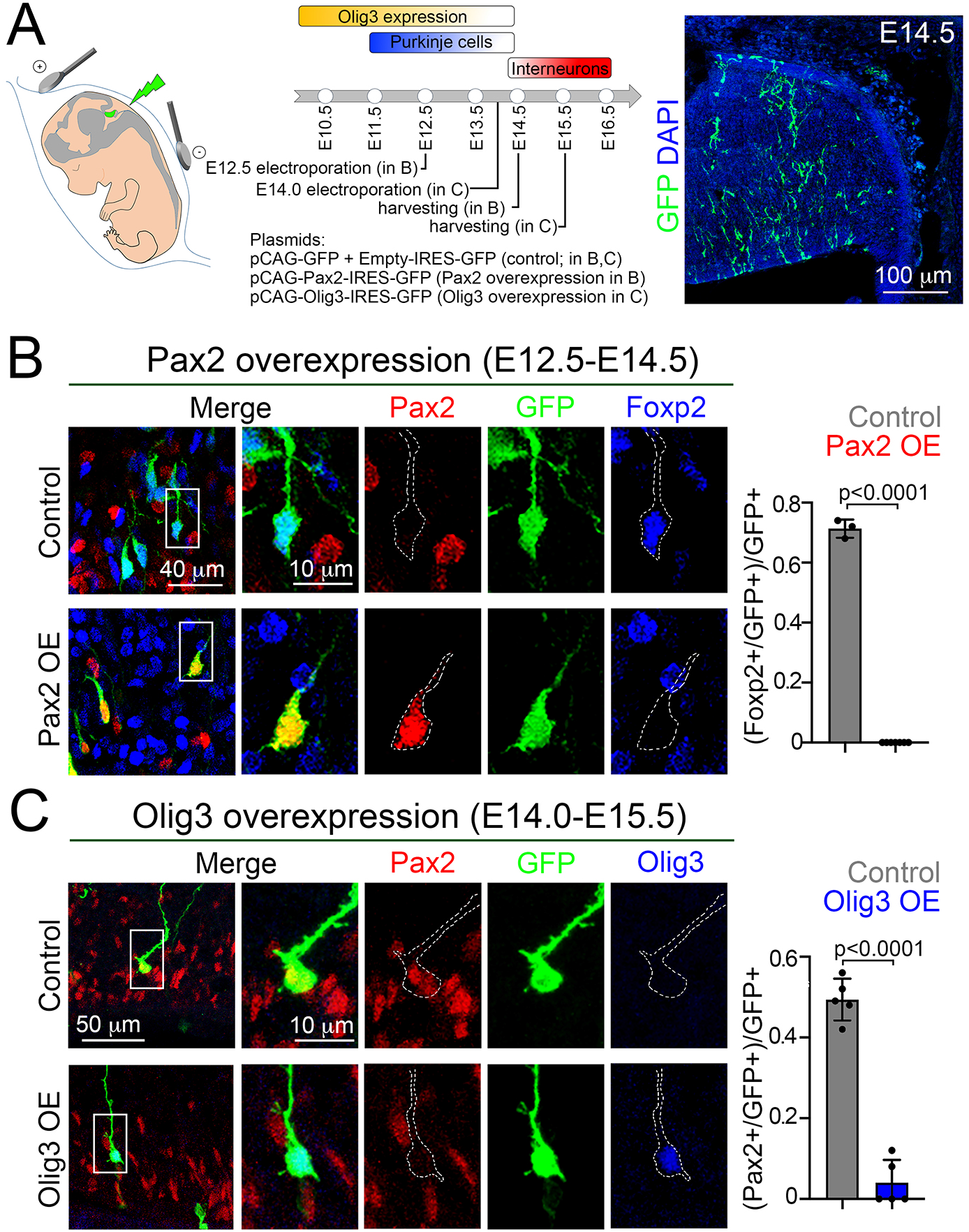
Pax2 imposes an inhibitory interneuron fate that is cell-autonomously suppressed by Olig3. (**A**) Strategy to force *Olig3* and *Pax2* expression in the ventricular zone of wildtype mouse embryos. Left, illustration of the electrode position required to target the ventricular zone. Middle top, schema illustrating the temporal expression of Olig3, and the generation of Purkinje cells and inhibitory interneurons. Middle bottom, electroporation of *Pax2* and *Olig3* expressing vectors was done at E12.5 and E14.0, respectively. Electroporated embryos were harvested at the indicated stages. Electroporated plasmids are shown. Right, a representative cerebellar section stained with GFP (green) and DAPI (blue) of an E14.5 mouse that was electroporated with control plasmids at E12.5. (**B**) Analysis of E14.5 wildtype mice that were electroporated at E12.5 with control (*pCAG-GFP + Empty-IRES-GFP*) or Pax2-overexpression (*pCAG-Pax2-IRES-GFP*) plasmids. Left, representative analyzed cells in the cerebellum of electroporated embryos that were stained against Pax2 (red), GFP (green) and Foxp2 (blue). Right, quantification of the proportion of GFP+ cells co-expressing Foxp2 in electroporated control (Pax2-) and Pax2-overexpressing (Pax2+) mice. (**C**) Analysis of E15.5 wildtype mice that were electroporated at E14.0 with control (*pCAG-GFP + Empty-IRES-GFP*) or Olig3-overexpression (*pCAG-Olig3-IRES-GFP*) plasmids. Left, representative analyzed cells in the cerebellum of electroporated embryos that were stained against Pax2 (red), GFP (green) and Olig3 (blue). Right, quantification of the proportion of GFP+ cells co-expressing Pax2 in electroporated control (Olig3-) and Olig3-overexpressing (Olig3+) mice. See Figure S6C for additional examples of electroporated cells. The mean and SD are plotted in all graphs, and the dots represent the mean of individual animals. Significance was determined using two-tailed t-tests, see Table S2. Photomicrographs were manually acquired using a Leica SPL confocal microscope.

To assess whether Olig3 cell-autonomously suppresses *Pax2* expression, we forced the ectopic expression of *Olig3* in the ventricular zone of wildtype mice at E14, a timepoint when *Olig3* expression is almost absent and Pax2+ cells initiate their specification. Electroporated *Olig3-IRES-GFP (Olig3* overexpressing) and *pCAG-GFP* + *Empty-IRES-GFP* (control) embryos were analyzed at E15.5. This showed that the proportion of GFP+ cells that co-expressed Pax2 was greatly reduced in the ventricular zone of Olig3-overexpressing embryos when compared to control electroporated mice (Figures 5C and S6C). Thus, expression of *Olig3* is sufficient to cell-autonomously suppress Pax2 expression. We conclude that during early development, Olig3 in the ventricular zone suppresses *Pax2* in newborn Purkinje cells to prevent their misspecification and secure their identity.

### Olig3 and Olig2 specify complementary Purkinje cell populations

Our analysis of bHLH factors expressed throughout cerebellar development showed that in addition to *Olig3* and *Ptf1a, Ascl1* and *Olig2* are also expressed in the ventricular zone during Purkinje cell generation (Figure 1A and Table S1). While ablation of *Ascl1* does not interfere with Purkinje cell development (Grimaldi et al., 2009; Sudarov et al., 2011), the exact role of Olig2 in the generation of GABAergic derivatives is unclear (Ju et al., 2016; Seto et al., 2014). In order to clarify the function of Olig2 we analyzed *Olig2* null mutant mice and found a reduction in Purkinje cell numbers and a modest increase in inhibitory interneurons (Figures S7A and S7B). We next carried out a long-term lineage tracing experiment using *Olig2^cre^* and *Tau^nLacZ^* alleles (*Olig2^cre/+^;Tau^nLacZ/+^* mice) to determine the contribution of Olig2 to cerebellar GABAergic neurons. This showed that while roughly half of the Foxp2+ Purkinje cell population had a history of *Olig2* expression, few inhibitory interneurons were generated from Olig2+ progenitors (Figures S7C and S7D). We thus conclude that the phenotypes of *Olig3* and *Olig2* mutant mice partially overlap (summarized in Figures 6A and S7D).

**Figure 6.**
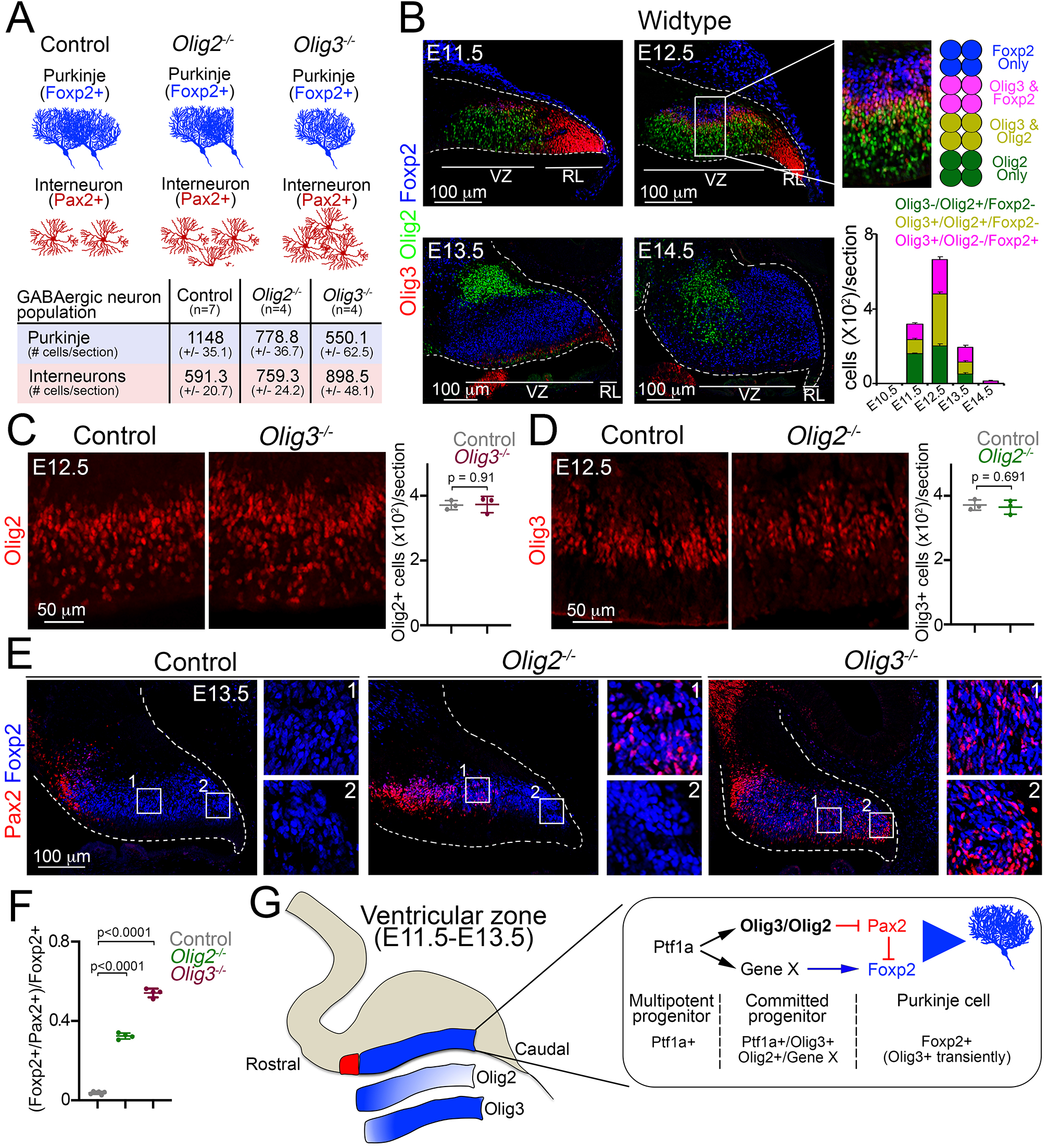
Complementary functions of Olig3 and Olig2 during Purkinje cell development. (**A**) Schema and quantification of the phenotypes observed in *Olig2* and *Olig3* mutant mice with respect to the development of GABAergic cerebellar neurons. See also Figure 3E, 3F (*Olig3* mutant analysis) and Figure S7A, B (*Olig2* mutant analysis). (**B**) Immunofluorescence characterization and quantification of Olig3+ (red), Olig2+ (green) and Foxp2+ (blue) cells in the ventricular zone at indicated embryonic stages. (**C**) Immunofluorescence characterization and quantification of Olig2+ (red) cells in the ventricular zone of *Olig3 (Olig3^-/-^*) mutant mice at E12.5. See also Figure S7E. (**D**) Immunofluorescence characterization and quantification of Olig3+ (red) cells in the ventricular zone of *Olig2 (Olig2^-/-^*) mutant mice at E12.5. See also Figure S7F. (**E**) Immunofluorescence comparison of Foxp2+ (blue) Purkinje cells and Pax2+ (red) inhibitory neurons in control versus *Olig2^-/-^* and *Olig3^-/-^* mutant mice at E13.5. Numbered boxed areas are displayed to the right of the main photographs. (**F**) Quantification of the proportion of Foxp2+ cells co-expressing Pax2 in control, *Olig2^-/-^* and *Olig3^-/-^* mutant mice at E13.5. (**G**) Schematic summary explaining the function of Olig3 and Olig2 in the ventricular zone during the specification of Purkinje cells. Induction of Olig3 and Olig2, in committed Ptf1a+ progenitor cells, curtails the expression of Pax2 to allow for the correct specification of Purkinje cells. Olig2 predominantly operates in the rostral ventricular zone, whereas Olig3 has a broader function and is transiently retained in newborn Purkinje cells. The suppression of Pax2 is critical for Purkinje cell development, as it can override the Purkinje cell differentiation program and instead impose an inhibitory interneuron identity. The mean and SD are plotted in all graphs, and the dots represent the mean of individual animals. Significance was determined using a one-way ANOVA followed by *post hoc* Tukey (in F) or two-tailed t-test (in C and D) analyses, see Table S2. Photomicrographs were acquired using the automatic tile scan modus (10% overlap between tiles) of the Zeiss spinning disk confocal microscope (in C-E) and the Zeiss LSM700 confocal microscope (in B).

Olig3 and Olig2 are known to specify non-overlapping neuron populations during the development of the hindbrain and spinal cord (Takebayashi et al., 2002; Takebayashi et al., 2000). To determine whether Olig3 and Olig2 mark complementary ventricular zone progenitor cells that specify distinct Foxp2+ Purkinje cells, we stained the cerebella of E10.5-E14.5 wildtype embryos with antibodies against these two factors. We observed that roughly 58% of Olig3+ cells in the ventricular zone co-expressed Olig2 but not Foxp2, while the remaining 42% of the Olig3+ cells co-expressed Foxp2 but not Olig2 (Figure 6B). Notably, we observed no Olig3-/Olig2+ cells that coexpressed Foxp2, illustrating that differentiated Purkinje cells retain *Olig3* but not *Olig2* expression.

We then asked whether ablation of *Olig3* might affect the expression of *Olig2* in ventricular zone progenitors, and vice versa we assessed whether mutation of *Olig2* might compromise the expression of *Olig3.* Mutation of *Olig3* did not affect the expression of *Olig2,* and neither did mutation of *Olig2* affect the expression of *Olig3* in progenitor cells of the ventricular zone (Figures 6C, 6D, S7E and S7F). These data demonstrate that the expression of *Olig3* and *Olig2* in ventricular zone progenitor cells is independent of the other factor. Next, we assessed whether mutation of *Olig2* might also de-repress Pax2 in newborn Purkinje cells in a similar manner as the ablation of *Olig3.* Indeed, we observed numerous Foxp2+/Pax2+ misspecified cells in *Olig2* mutant animals at E13.5 (Figure 6E, quantified in Figure 6F), but unlike in *Olig3* mutant embryos, these cells were only located in the rostral-most part of the ventricular zone (compare insets in Figure 6E). These data demonstrate that Olig2 specifically suppresses *Pax2* in rostrally generated Purkinje cells, while Olig3 has a broader function in the suppression of *Pax2* in most of the Purkinje cell population. We therefore conclude that Olig3 and Olig2 complementarily contribute to the correct specification of Purkinje cells by curtailing the expression of *Pax2* (schematically displayed in Figure 6G).

## Discussion

bHLH transcription factors are highly conserved in evolution and function as principal regulators of cell differentiation and neuronal specification (Atchley and Fitch, 1997; Baker and Brown, 2018; Ben-Arie et al., 2000; Bertrand et al., 2002; Dennis et al., 2019; Dokucu et al., 1996; Jones, 2004; Sommer et al., 1996). In this study, we sought to identify bHLH factors that regulate the specification of distinct cerebellar neuron types. We report here that Olig3 is a key player in cerebellar development and the generation of its earliest neuronal derivatives. Ablation of *Olig3* results in pronounced cerebellar hypoplasia at birth and the massive loss of DCN neurons, EGL cells including their granular cell derivatives, and Purkinje cells. These deficits are accompanied by an increase in the number of inhibitory interneurons. We show that Olig3 cell-autonomously suppresses the development of inhibitory interneurons by curtailing the expression of *Pax2* both in the rhombic lip and ventricular zone. We also demonstrate that Pax2 acts as an effective selector gene that imposes an inhibitory interneuron identity. In addition, we show that Olig3 and its close family member Olig2 specify complementary Purkinje cell populations. Thus, our work defines Olig3 as a master regulator of cerebellar development and is required for the specification of the earliest set of cerebellar neuron types.

Here, we show that Olig3 is critically involved in the generation of EGL cells as well as DCN neurons (Ben-Arie et al., 1997; Fink et al., 2006; Gazit et al., 2004; Machold and Fishell, 2005; Machold et al., 2011). Earlier studies revealed that these rhombic lip derivatives depend on Atoh1 for their development, as loss of Atoh1 results in the severe reduction of EGL cells and impairs the development of DCN neurons (Ben-Arie et al., 1997; Jensen et al., 2004; Wang et al., 2005; Yamada et al., 2014). Our long-term lineage tracing studies demonstrated that most EGL and DCN cells have a history of *Olig3* expression, ablation of which massively reduced their cell numbers. In the early rhombic lip (E10.5-E13.5) we found that most proliferative progenitor cells (Sox2+/BrdU+) co-expressed Olig3 and a third of them co-expressed Atoh1 (Olig3+/Atoh1+ cells). This temporal window overlaps with the generation of DCN cells (Fink et al., 2006; Sekerkova et al., 2004; Wang et al., 2005; Yamada et al., 2014), which are the most reduced neuron type in *Olig3* mutant mice (this study). Thus, Olig3 is essential for DCN neuron development. Ablation of *Olig3* reduced the number of BrdU+ (proliferative) progenitor cells in the rhombic lip, and consequently decreased the number of Atoh1+ cells. This impairment led to smaller numbers of EGL cells and, therefore, to fewer differentiated granular cells. The severe loss of EGL cells and their granular cell derivatives seems to largely account for the pronounced cerebellar hypoplasia observed in *Olig3* mutant mice, and has also been observed after the loss of EGL cells in other studies(Ben-Arie et al., 1997). During the development of rhombomeres 2 to 7, there exists a dorsal progenitor domain (called dA1) that also co-expresses Olig3 and Atoh1 (Hernandez-Miranda et al., 2017a; Liu et al., 2008; Storm et al., 2009). This domain generates the mossy fiber precerebellar (pontine, lateral reticular, external cuneate) nuclei. Like in the rhombic lip, ablation of *Olig3* greatly reduces the number of Atoh1+ cells in this area and their derivatives (Liu et al., 2008; Storm et al., 2009). This phenomenon also occurs in the spinal cord when *Olig3* is ablated (Muller et al., 2005). Thus, Olig3 has a conserved function in the proliferation of Atoh1+ progenitor cells.

We also show that the ablation of *Olig3* results in the development of supernumerary inhibitory interneurons. Both Purkinje cells and inhibitory interneurons depend on Ptf1a for their development (Hashimoto and Mikoshiba, 2003; Hoshino et al., 2005; Leto et al., 2006; Yamada et al., 2014). We predominantly found expression of Olig3 in the ventricular zone between E11.5 and E13.5, the temporal window during which Purkinje cells are specified. In the ventricular zone, about half of the Olig3+ cells co-express Ptf1a, while the rest co-express the Purkinje cell marker Foxp2. This shows that Olig3 expression is initiated in progenitors and transiently retained in newborn Purkinje cells. Ablation of *Olig3* neither impaired the number of Ptf1a+ cells nor their proliferation. Strikingly, around half of newborn Purkinje cells erroneously co-expressed Pax2 in *Olig3* mutant mice at E13.5, illustrating that ablation of *Olig3* misspecifies newborn Purkinje cells. In *Olig3* mutant mice, the number of misspecified cells declined over time and became rare by P0. This decline correlated with a parallel increase in the number of inhibitory interneurons. Thus, the primary function of Olig3 in the ventricular zone is to secure the development of Purkinje cells by cell-autonomously suppressing an alternative program that specifies inhibitory interneurons. In this context, our functional data demonstrate that forced expression of *Olig3,* during the temporal generation of inhibitory interneurons, is sufficient to curtail *Pax2* expression. Furthermore, our functional data demonstrate that Pax2 acts as an effective selector gene in the specification of inhibitory interneurons and can override *Foxp2* expression. The suppression of *Pax2* by Olig3 is not limited to ventricular zone progenitor cells as we also observed expression of Pax2 in GFP+ cells in the rhombic lip of *Olig3^GFP/GFP^* mutant mice. In agreement with our findings, it was previously shown that supernumerary inhibitory neurons become specified at the expense of excitatory neurons in the hindbrain and spinal cord of *Olig3* mutant mice (Muller et al., 2005; Storm et al., 2009; Zechner et al., 2007). Interestingly, there is a unique progenitor domain in rhombomere 7 (called dA4) that co-expresses Olig3 and Ptf1a, which generates the precerebellar climbing fiber neurons of the inferior olive (reviewed in (Hernandez-Miranda et al., 2017a). In the absence of Olig3, inferior olive neurons and many spinal cord excitatory neurons seem to change their fate and erroneously adopt an inhibitory interneuron identity (Liu et al., 2008; Muller et al., 2005; Storm et al., 2009). This suggests that inhibitory interneurons are the default neuronal type generated from the cerebellum, brainstem and spinal cord. Thus, Olig3 has a conserved function in the suppression of inhibitory interneuron specification during development.

Based on short-term lineage tracing experiments, Seto and colleagues (2014) postulated a “temporal identity transition” model in which Olig2+ Purkinje cell progenitors transition into inhibitory interneuron progenitors (Seto et al., 2014). From this model, one would expect that inhibitory interneurons would have a history of *Olig2* expression. In keeping with observations made by Ju et al. (Ju et al., 2016), our long-term lineage tracing experiments using *Olig2^cre^* and *Olig3^creERT2^* mice showed that Pax2+ inhibitory interneurons rarely have a history of *Olig2* or *Olig3* expression. This casts doubt on the “temporal identity transition” model as both factors are abundantly expressed in Ptf1a+ progenitors during the specification of Purkinje cells (this work and Seto et al., 2014). Our data unambiguously show that neither Olig2 nor Olig3 control the transition of early (Purkinje) to late (inhibitory interneuron) ventricular zone progenitor cells. Rather, our work demonstrates that these factors are essential for the correct specification of Purkinje cells by curtailing an inhibitory interneuron transcriptional program.

Development of the central nervous system is characterized by molecular “grids” of combinatorial transcription factor expression that single out distinct progenitor domains. It is from here that the enormous diversity of neuron types is generated (reviewed in (Alaynick et al., 2011; Hernandez-Miranda et al., 2017a; Hernandez-Miranda et al., 2010; Jessell, 2000). Here we show that cerebellar DCN neurons and internal granular cells develop from Olig3+/Atoh1 + rhombic lip progenitor cells, whereas Purkinje cells derive from Olig3+/Ptf1a+ ventricular zone progenitors. In the mature cerebellum, DCN neurons and granular cells receive input from brainstem precerebellar mossy fiber neurons that originate from progenitor cells that co-express Olig3 and Atoh1, whilst Purkinje cells receive input from climbing fiber neurons that emerge from progenitors that co-express Olig3 and Ptf1a (Liu et al., 2008; Storm et al., 2009). The question of how these progenitor cells, located at such distant positions, acquire similar molecular signatures to specify both targets and inputs that in turn form functional cerebellar circuits remains to be elucidated.

## Materials and Methods

### Animals

All animal experimental procedures were done in accordance to the guidance and policies of the Charite Universitatsmedizin, Berlin, Germany; Max-Delbrück-Center for Molecular Medicine, Berlin, Germany; and the Institute of Neuroscience, Lobachevsky University of Nizhny Novgorod, Russian Federation. Mouse strains used for this study were: *Olig3creERT2(Storm* et al., 2009), *Olig3GFP(Muller* et al., 2005), *TaunLacZ* (Hippenmeyer et al., 2005), *Ail14* (Madisen et al., 2010) and *Olig2cre*(Dessaud et al., 2007). All strains were maintained in a mixed genetic background.

For tamoxifen treatment, pregnant dams were treated with tamoxifen (Sigma-Aldrich; 20 mg/ml dissolved in sunflower oil) as described previously(Hernandez-Miranda et al., 2017b; Storm et al., 2009). Tamoxifen delays labor in rodents and humans(Lizen et al., 2015); therefore, offspring from tamoxifen-treated dams were delivered by cesarian section at E19.

### Histology and cell quantifications

Immunofluorescence and tissue processing were performed as previously described(Hernandez-Miranda et al., 2011). Briefly, mouse tissue (E10.5–P0) was fixed in 4% paraformaldehyde (PFA), made in phosphate buffered saline (PBS), for 3 hours at 4°C. After fixation, brains were cryoprotected in 30% sucrose in PBS, embedded and frozen in Tissue-Tek OCT (Sakura Finetek), and sectioned at 20 μm using a cryostat. Sections were washed in PBS and blocked in PBS containing 5% normal goat serum (Sigma-Aldrich) (v/v) and 0.1% Triton X-100 (v/v) (Sigma-Aldrich) at room temperature for 2 hours. They were subsequently incubated in primary antibodies at room temperature overnight. After incubation in primary antibodies, sections were washed in PBS and then incubated in secondary antibodies for 2 hours at room temperature. The following primary antibodies were used in this study: rabbit anti-Atoh1 (1:10000, kindly provided by Thomas Jessell), chicken anti-bgal (1:1000, Abcam, ab9361), rat anti-BrdU (1:2000, Abcam, ab6326), goat anti-Brn2 (1:1000, Abcam, ab101726), rabbit anti-Caspase3 (1:1000, R&D Systems, AF835), goat anti-Foxp2 (1:1000, Abcam, ab58599), rabbit anti-Foxp2 (1:1000, Abcam, ab16046), chicken anti-GFP (1:500, Abcam, ab13970), rabbit anti-GFP (1:500, Abcam, ab290), rat anti-GFP (1:2000, Nacalai Tesque, GF090R), rabbit anti-Olig2 (1:1000, Merck Millipore, AB9610), guinea pig anti-Olig3 (1:5000, ref. (Muller et al., 2005)), rabbit anti-Pax2 (1:1000, Abcam, EP3251), rabbit anti-Pax6 (1:1000, Merck Millipore, AB2237), rabbit anti-Ptf1a (1:5000, kindly provided by Jane Johnson), rabbit anti-RFP (1:500, Rockland, 600-401-379-RTU), rabbit anti-Sox2 (1:1000, Abcam, ab97959), rabbit anti-Tbr1 (1:1000, Abcam, EPR8138) and rabbit anti-Tbr2 (1:1000, Merck Millipore, AB15894). Species-specific Cy2-, Cy3-, and Cy5-conjugated secondary antibodies were obtained from Jackson ImmunoResearch and used 1:1000. For a 45-minute BrdU pulse labeling, BrdU (Sigma-Aldrich) was diluted to a concentration of 16 mg/ml in saline solution and injected intraperitoneally.

Cell quantifications were performed in a non-blind manner on non-consecutive 20μm-thick brain sections encompassing the complete lateral-medial cerebellar axis. On average six to ten sections per animal were used for quantifications. E12.5 whole-mount embryos were analysed for β-gal activity with X-gal (0.6 mg/ml; Merck Millipore, B4252) in PBS buffer containing 4 mM potassium ferricyanide, 4 mM potassium ferrocyanide, 0.02% NP-40 and 2 mM MgCl2 as previously described(Comai and Tajbakhsh, 2014). For the estimation of the cerebellar volume and area, consecutive 20-μm-thick sagittal sections were collected encompassing the whole cerebellum and stained with Nissl. Roughly 32-35 sections of the cerebellum were obtained per animal (four animals/genotype). The area of every section was measured using ImageJ; NIH, version 1.34n. Estimation of the total volume of the cerebellum was obtained by application of Cavalieri’s method(West, 2012). Fluorescence images were acquired using: i) a Zeiss LSM 700 confocal microscope using the automatic tile scan modus (10% overlap between tiles) and assembled using ZEN2012, ii) a Zeiss spinning disk confocal microscope using the automatic tile scan modus (10% overlap between tiles) and assembled using ZEN2012, and iii) a Leica SPL confocal microscope. Photographs obtained with the Leica SPL confocal microscope were manually acquired and these were assembled using Image J. Unless otherwise specified all photomicrographs were acquired in a non-blind manner.

### Brain clearing, light-sheet microscopy and analysis

Brains were cleared using the CUBIC protocol(Susaki et al., 2015). Briefly, brains were dissected and fixed overnight at 4°C in 4% paraformaldehyde made in PBS. After washing overnight in PBS, lipids were removed using Reagent-1 (25% urea, 25% Quadrol, 15% Triton X-100, 35% dH_2_O) at 37°C until brains were transparent (4 days). The brains were then washed overnight at 4°C in PBS to remove Reagent-1 and then placed into Reagent-2 (25% urea, 50% sucrose, 10% triethanolamine, 15% dH_2_O) at 37°C for refractive index matching (3 days). Once the brains were cleared they were imaged using a Zeiss Lightsheet Z.1 microscope. 3D reconstruction, photos and videos were created with arivis Vision4D.

### *In utero* electroporation

In utero electroporation was performed as previously described(Saito and Nakatsuji, 2001). Briefly, DNA plasmids were mixed with Fast Green and injected into the fourth ventricle of embryonic brains from outside the uterus with a glass micropipette. Holding the embryo in utero with forceps-type electrodes (NEPA GENE), 50 ms of 40 V electronic pulses were delivered five times at intervals of 950 ms with a square electroporator (Nepa Gene, CUY21). The plasmids used in this study were: *pCAG-GFP* (control), *pCAG-Empty-IRES-GFP* (control), *pCAG-Olig3-IRES-GFP* (Olig3 overexpression) and *pCAG-Pax2-IRES-GFP.* Primers used to clone the mouse *Olig3* (NM_053008.3) gene in *pCAG-Empty-IRES-GFP* were: forward: ATGAATTCTGATTCGAGC and reverse: TTAAACCTTATCGTCGTC. Primers used to clone mouse *Pax2* (NM_011037.5) gene in *pCAG-Empty-IRES-GFP* were: forward: ATGGATATGCACTGCAAAGCAG and reverse: GTGGCGGTCATAGGCAGC. The electroporated plasmid DNA mixtures were as follows: i) for the control experiment, *pCAG-GFP* (0.5 mg ml^-1^) + *pCAG-Empty-IRES-GFP* (0.5 mg ml^-1^); ii) for the *Olig3* overexpression experiment*, pCAG-Olig3-IRES-GFP* (0.5 mg ml^-1^); and iii) for the *Pax2* overexpression experiment, *pCAG-Pax2-IRES-GFP* (0.5 mg ml^-1^).

### Statistics

Statistical analyses were performed using Prism 8 (GraphPad). Data are plotted in scatter dot plots or column dot plots with means and standard deviations (SD) displayed. The statistical significance between group means was tested by one-way ANOVA, followed by Tukey’s *post hoc* test (for multiple-comparison tests), or two-tailed t-test (for pair comparison tests). Degrees of Freedom as well as F and t values are provided in Table S2.

## References

Alaynick, W.A., Jessell, T.M., and Pfaff, S.L. (2011). SnapShot: spinal cord development. Cell 146, 178–178 e171.

Alder, J., Cho, N.K., and Hatten, M.E. (1996). Embryonic precursor cells from the rhombic lip are specified to a cerebellar granule neuron identity. Neuron 17, 389–399.

Atchley, W.R., and Fitch, W.M. (1997). A natural classification of the basic helixloop-helix class of transcription factors. Proc Natl Acad Sci U S A 94, 5172–5176.

Baker, N.E., and Brown, N.L. (2018). All in the family: proneural bHLH genes and neuronal diversity. Development 145.

Ben-Arie, N., Bellen, H.J., Armstrong, D.L., McCall, A.E., Gordadze, P.R., Guo, Q., Matzuk, M.M., and Zoghbi, H.Y. (1997). Math1 is essential for genesis of cerebellar granule neurons. Nature 390, 169–172.

Ben-Arie, N., Hassan, B.A., Bermingham, N.A., Malicki, D.M., Armstrong, D., Matzuk, M., Bellen, H.J., and Zoghbi, H.Y. (2000). Functional conservation of atonal and Math1 in the CNS and PNS. Development 127, 1039–1048.

Bertrand, N., Castro, D.S., and Guillemot, F. (2002). Proneural genes and the specification of neural cell types. Nat Rev Neurosci 3, 517–530.

Butts, T., Green, M.J., and Wingate, R.J. (2014). Development of the cerebellum: simple steps to make a ‘little brain’. Development 141, 4031–4041.

Chizhikov, V.V., Lindgren, A.G., Currle, D.S., Rose, M.F., Monuki, E.S., and Millen, K.J. (2006). The roof plate regulates cerebellar cell-type specification and proliferation. Development 133, 2793–2804.

Comai, G., and Tajbakhsh, S. (2014). Molecular and cellular regulation of skeletal myogenesis. Curr Top Dev Biol 110, 1–73.

Dennis, D.J., Han, S., and Schuurmans, C. (2019). bHLH transcription factors in neural development, disease, and reprogramming. Brain Res 1705, 48–65.

Dessaud, E., Yang, L.L., Hill, K., Cox, B., Ulloa, F., Ribeiro, A., Mynett, A., Novitch, B.G., and Briscoe, J. (2007). Interpretation of the sonic hedgehog morphogen gradient by a temporal adaptation mechanism. Nature 450, 717–720.

Dokucu, M.E., Zipursky, S.L., and Cagan, R.L. (1996). Atonal, rough and the resolution of proneural clusters in the developing Drosophila retina. Development 122, 4139–4147.

Englund, C., Kowalczyk, T., Daza, R.A., Dagan, A., Lau, C., Rose, M.F., and Hevner, R.F. (2006). Unipolar brush cells of the cerebellum are produced in the rhombic lip and migrate through developing white matter. J Neurosci 26, 9184–9195.

Fink, A.J., Englund, C., Daza, R.A., Pham, D., Lau, C., Nivison, M., Kowalczyk, T., and Hevner, R.F. (2006). Development of the deep cerebellar nuclei: transcription factors and cell migration from the rhombic lip. J Neurosci 26, 3066–3076.

Gazit, R., Krizhanovsky, V., and Ben-Arie, N. (2004). Math1 controls cerebellar granule cell differentiation by regulating multiple components of the Notch signaling pathway. Development 131, 903–913.

Grimaldi, P., Parras, C., Guillemot, F., Rossi, F., and Wassef, M. (2009). Origins and control of the differentiation of inhibitory interneurons and glia in the cerebellum. Dev Biol 328, 422–433.

Hallonet, M.E., Teillet, M.A., and Le Douarin, N.M. (1990). A new approach to the development of the cerebellum provided by the quail-chick marker system. Development 108, 19–31.

Hashimoto, M., and Mikoshiba, K. (2003). Mediolateral compartmentalization of the cerebellum is determined on the “birth date” of Purkinje cells. J Neurosci 23, 11342–11351.

Hernandez-Miranda, L.R., Cariboni, A., Faux, C., Ruhrberg, C., Cho, J.H., Cloutier, J.F., Eickholt, B.J., Parnavelas, J.G., and Andrews, W.D. (2011). Robo1 regulates semaphorin signaling to guide the migration of cortical interneurons through the ventral forebrain. J Neurosci 31, 6174–6187.

Hernandez-Miranda, L.R., Muller, T., and Birchmeier, C. (2017a). The dorsal spinal cord and hindbrain: From developmental mechanisms to functional circuits. Dev Biol 432, 34–42.

Hernandez-Miranda, L.R., Parnavelas, J.G., and Chiara, F. (2010). Molecules and mechanisms involved in the generation and migration of cortical interneurons. ASN Neuro 2, e00031.

Hernandez-Miranda, L.R., Ruffault, P.L., Bouvier, J.C., Murray, A.J., Morin-Surun, M.P., Zampieri, N., Cholewa-Waclaw, J.B., Ey, E., Brunet, J.F., Champagnat, J., et al. (2017b). Genetic identification of a hindbrain nucleus essential for innate vocalization. Proc Natl Acad Sci U S A 114, 8095–8100.

Hippenmeyer, S., Vrieseling, E., Sigrist, M., Portmann, T., Laengle, C., Ladle, D.R., and Arber, S. (2005). A developmental switch in the response of DRG neurons to ETS transcription factor signaling. PLoS Biol 3, e159.

Hoshino, M., Nakamura, S., Mori, K., Kawauchi, T., Terao, M., Nishimura, Y.V., Fukuda, A., Fuse, T., Matsuo, N., Sone, M., et al. (2005). Ptf1a, a bHLH transcriptional gene, defines GABAergic neuronal fates in cerebellum. Neuron 47, 201–213.

Imayoshi, I., and Kageyama, R. (2014). bHLH factors in self-renewal, multipotency, and fate choice of neural progenitor cells. Neuron 82, 9–23.

Jensen, P., Smeyne, R., and Goldowitz, D. (2004). Analysis of cerebellar development in math1 null embryos and chimeras. J Neurosci 24, 2202–2211.

Jessell, T.M. (2000). Neuronal specification in the spinal cord: inductive signals and transcriptional codes. Nat Rev Genet 1, 20–29.

Jones, S. (2004). An overview of the basic helix-loop-helix proteins. Genome Biol 5, 226.

Ju, J., Liu, Q., Zhang, Y., Liu, Y., Jiang, M., Zhang, L., He, X., Peng, C., Zheng, T., Lu, Q.R., et al. (2016). Olig2 regulates Purkinje cell generation in the early developing mouse cerebellum. Sci Rep 6, 30711.

Leto, K., Carletti, B., Williams, I.M., Magrassi, L., and Rossi, F. (2006). Different types of cerebellar GABAergic interneurons originate from a common pool of multipotent progenitor cells. J Neurosci 26, 11682–11694.

Liu, Z., Li, H., Hu, X., Yu, L., Liu, H., Han, R., Colella, R., Mower, G.D., Chen, Y., and Qiu, M. (2008). Control of precerebellar neuron development by Olig3 bHLH transcription factor. J Neurosci 28, 10124–10133.

Liu, Z.R., Shi, M., Hu, Z.L., Zheng, M.H., Du, F., Zhao, G., and Ding, Y.Q. (2010). A refined map of early gene expression in the dorsal rhombomere 1 of mouse embryos. Brain Res Bull 82, 74–82.

Lizen, B., Claus, M., Jeannotte, L., Rijli, F.M., and Gofflot, F. (2015). Perinatal induction of Cre recombination with tamoxifen. Transgenic Res 24, 1065–1077.

Machold, R., and Fishell, G. (2005). Math1 is expressed in temporally discrete pools of cerebellar rhombic-lip neural progenitors. Neuron 48, 17–24.

Machold, R., Klein, C., and Fishell, G. (2011). Genes expressed in Atoh1 neuronal lineages arising from the r1/isthmus rhombic lip. Gene Expr Patterns 11, 349–359.

Madisen, L., Zwingman, T.A., Sunkin, S.M., Oh, S.W., Zariwala, H.A., Gu, H., Ng, L.L., Palmiter, R.D., Hawrylycz, M.J., Jones, A.R., et al. (2010). A robust and high-throughput Cre reporting and characterization system for the whole mouse brain. Nat Neurosci 13, 133–140.

Maricich, S.M., and Herrup, K. (1999). Pax-2 expression defines a subset of GABAergic interneurons and their precursors in the developing murine cerebellum. J Neurobiol 41, 281–294.

Mattar, P., Langevin, L.M., Markham, K., Klenin, N., Shivji, S., Zinyk, D., and Schuurmans, C. (2008). Basic helix-loop-helix transcription factors cooperate to specify a cortical projection neuron identity. Mol Cell Biol 28, 1456–1469.

Millen, K.J., Steshina, E.Y., Iskusnykh, I.Y., and Chizhikov, V.V. (2014). Transformation of the cerebellum into more ventral brainstem fates causes cerebellar agenesis in the absence of Ptf1a function. Proc Natl Acad Sci U S A 111, E1777–1786.

Millet, S., Bloch-Gallego, E., Simeone, A., and Alvarado-Mallart, R.M. (1996). The caudal limit of Otx2 gene expression as a marker of the midbrain/hindbrain boundary: a study using in situ hybridisation and chick/quail homotopic grafts. Development 122, 3785–3797.

Morales, D., and Hatten, M.E. (2006). Molecular markers of neuronal progenitors in the embryonic cerebellar anlage. J Neurosci 26, 12226–12236.

Muller, T., Anlag, K., Wildner, H., Britsch, S., Treier, M., and Birchmeier, C. (2005). The bHLH factor Olig3 coordinates the specification of dorsal neurons in the spinal cord. Genes Dev 19, 733–743.

Ross, S.E., Greenberg, M.E., and Stiles, C.D. (2003). Basic helix-loop-helix factors in cortical development. Neuron 39, 13–25.

Saito, T., and Nakatsuji, N. (2001). Efficient gene transfer into the embryonic mouse brain using in vivo electroporation. Dev Biol 240, 237–246.

Sekerkova, G., Ilijic, E., and Mugnaini, E. (2004). Time of origin of unipolar brush cells in the rat cerebellum as observed by prenatal bromodeoxyuridine labeling. Neuroscience 127, 845–858.

Sellick, G.S., Barker, K.T., Stolte-Dijkstra, I., Fleischmann, C., Coleman, R.J., Garrett, C., Gloyn, A.L., Edghill, E.L., Hattersley, A.T., Wellauer, P.K., et al. (2004). Mutations in PTF1A cause pancreatic and cerebellar agenesis. Nat Genet 36, 1301–1305.

Seto, Y., Nakatani, T., Masuyama, N., Taya, S., Kumai, M., Minaki, Y., Hamaguchi, A., Inoue, Y.U., Inoue, T., Miyashita, S., et al. (2014). Temporal identity transition from Purkinje cell progenitors to GABAergic interneuron progenitors in the cerebellum. Nat Commun 5, 3337.

Shiraishi, A., Muguruma, K., and Sasai, Y. (2017). Generation of thalamic neurons from mouse embryonic stem cells. Development 144, 1211–1220.

Sommer, L., Ma, Q., and Anderson, D.J. (1996). neurogenins, a novel family of atonal-related bHLH transcription factors, are putative mammalian neuronal determination genes that reveal progenitor cell heterogeneity in the developing CNS and PNS. Mol Cell Neurosci 8, 221–241.

Storm, R., Cholewa-Waclaw, J., Reuter, K., Brohl, D., Sieber, M., Treier, M., Muller, T., and Birchmeier, C. (2009). The bHLH transcription factor Olig3 marks the dorsal neuroepithelium of the hindbrain and is essential for the development of brainstem nuclei. Development 136, 295–305.

Sudarov, A., Turnbull, R.K., Kim, E.J., Lebel-Potter, M., Guillemot, F., and Joyner, A.L. (2011). Ascl1 genetics reveals insights into cerebellum local circuit assembly. J Neurosci 31, 11055–11069.

Susaki, E.A., Tainaka, K., Perrin, D., Yukinaga, H., Kuno, A., and Ueda, H.R. (2015). Advanced CUBIC protocols for whole-brain and whole-body clearing and imaging. Nat Protoc 10, 1709–1727.

Takebayashi, H., Ohtsuki, T., Uchida, T., Kawamoto, S., Okubo, K., Ikenaka, K., Takeichi, M., Chisaka, O., and Nabeshima, Y. (2002). Non-overlapping expression of Olig3 and Olig2 in the embryonic neural tube. Mech Dev 113, 169–174.

Takebayashi, H., Yoshida, S., Sugimori, M., Kosako, H., Kominami, R., Nakafuku, M., and Nabeshima, Y. (2000). Dynamic expression of basic helixloop-helix Olig family members: implication of Olig2 in neuron and oligodendrocyte differentiation and identification of a new member, Olig3. Mech Dev 99, 143–148.

Vue, T.Y., Aaker, J., Taniguchi, A., Kazemzadeh, C., Skidmore, J.M., Martin, D.M., Martin, J.F., Treier, M., and Nakagawa, Y. (2007). Characterization of progenitor domains in the developing mouse thalamus. J Comp Neurol 505, 73–91.

Wang, V.Y., Rose, M.F., and Zoghbi, H.Y. (2005). Math1 expression redefines the rhombic lip derivatives and reveals novel lineages within the brainstem and cerebellum. Neuron 48, 31–43.

West, M.J. (2012). Estimating volume in biological structures. Cold Spring Harb Protoc 2012, 1129–1139.

Wingate, R.J., and Hatten, M.E. (1999). The role of the rhombic lip in avian cerebellum development. Development 126, 4395–4404.

Yamada, M., Seto, Y., Taya, S., Owa, T., Inoue, Y.U., Inoue, T., Kawaguchi, Y., Nabeshima, Y., and Hoshino, M. (2014). Specification of spatial identities of cerebellar neuron progenitors by ptf1a and atoh1 for proper production of GABAergic and glutamatergic neurons. J Neurosci 34, 4786–4800.

Yeung, J., Ha, T.J., Swanson, D.J., and Goldowitz, D. (2016). A Novel and Multivalent Role of Pax6 in Cerebellar Development. J Neurosci 36, 9057–9069.

Zechner, D., Muller, T., Wende, H., Walther, I., Taketo, M.M., Crenshaw, E.B., 3rd, Treier, M., Birchmeier, W., and Birchmeier, C. (2007). Bmp and Wnt/beta-catenin signals control expression of the transcription factor Olig3 and the specification of spinal cord neurons. Dev Biol 303, 181–190.

Zervas, M., Millet, S., Ahn, S., and Joyner, A.L. (2004). Cell behaviors and genetic lineages of the mesencephalon and rhombomere 1. Neuron 43, 345–357.

