## Supplementary Info for "Olig3 acts as a master regulator of cerebellar development"

#### Supplementary Information

#### Supplementary Figure Legends

**Supplementary Figure 1. Characterization of Olig3 expression during cerebellar development.**

(**A**) Sagittal wildtype cerebellar sections stained with antibodies against Olig3 (red) and DAPI (blue) between E10.5 to E14.5. The ventricular zone (VZ) and rhombic lip (RL) are indicated.

(**B, C**) Immunofluorescence analysis and quantification of proliferative BrdU+ (green) cells co-expressing Olig3 (red) in the rhombic lip (B) and the ventricular zone (C) between E10.5 to E14.5.

The mean and SD are plotted in all graphs, and the dots represent the mean of individual animals. Photomicrographs were acquired using the automatic tile scan modus (10% overlap between tiles) of the Zeiss LSM700 confocal microscope.

**Supplementary Figure 2. Lineage-tracing of Olig3-derived cells.**

(**A**) Strategy to label Olig3-derived cells using a tamoxifen-dependent cre recombinase driven by *Olig3* and the indicator allele Ai14 ($Olig3^{creERT2/+};Ai14^{+/-}$ mice). Upon tamoxifen (TAM) treatment, a stop cassette upstream of the Tomato coding sequence is removed. Recombined cells are visualized by Tomato fluorescence (red). (**A', A''**) A whole-mount three-dimensional reconstruction of an $Olig3^{creERT2/+};Ai14^{+/-}$ brain that was recombined at E10.5 and imaged at E19. DAPI (white) was used as a counterstain (A'). Note the distribution of Tomato+ cells across the cerebellum and DCN nuclei (asterisks in A''). Other derivatives of Olig3+ progenitors, such as the thalamus (Th) and its projections to the cortex (Cx), the pontine nuclei (Pn) and their projections (white arrowhead) or the inferior olive (Io) and its projections (yellow arrowhead) in the medulla oblongata (Mo) are also marked with Tomato.

(**B**) Strategy to label Olig3-derived neurons using $Olig3^{creERT2}$ and the indicator allele Tau nuclear-LacZ ($Olig3^{creERT2/+};Tau^{nLacZ/+}$ mice). (**B'**) To visualize Olig3 derivatives throughout development, one dose of tamoxifen was given to pregnant dams (note that mice only received tamoxifen on one day during development). Recombined animals were analyzed at E19. (**B''**) Lateral view of an E12.5 *Olig3*$^{creERT2/+}$;*Tau*$^{nLacZ/+}$ embryo that was recombined at E10.5. Note the broad expression of βgal, visualized using X-Gal staining, in the entire cerebellar anlage (arrowhead).

(**C**) Left, a sagittal cerebellar section, of an E19 *Olig3*$^{creERT2/+}$;*Tau*$^{nLacZ/+}$ mouse that was recombined with tamoxifen at E10.5, and stained against βgal (red) and DAPI (blue). Note that the external granular cell layer (arrows) does not express βgal (see text). Right, quantification of βgal+ cells in the cerebellum of E19 *Olig3*$^{creERT2/+}$;*Tau*$^{nLacZ/+}$ mice that were recombined at different embryonic stages (E10.5-E13.5).

(**D, E**) *Olig3*$^{creERT2/+}$;*Tau*$^{nLacZ/+}$ mice were recombined with tamoxifen at different embryonic stages and analyzed at E19. Sagittal sections from these mice were stained against βgal (red) and markers for dentatus/interpositus DCN neurons (Brn2, blue in D) and unipolar brush cells (Tbr2, blue in E). Double-positive (βgal+/marker+) cells were quantified at E19.

The mean and SD are plotted in all graphs, and the dots represent the mean of individual animals. Significance was obtained using one-way ANOVA followed by *post hoc* Tukey's test, see Table S2. Photomicrographs were acquired using the automatic tile scan modus (10% overlap between tiles) of the Zeiss LSM700 confocal microscope.

**Supplementary Figure 3. Hypoplasia and loss of defined neurons in the cerebellum of *Olig3* mutant mice.**

(**A**) Estimation of cerebellar volume in newborn control (*Olig3*$^{+/-}$) and *Olig3* mutant (*Olig3*$^{-/-}$) mice. The solid lines represent the mean of analyzed animals and the shaded areas the SD (n=4 mice/genotype), see also Figure 3A.

(**B, C**) Immunofluorescence characterization and quantification of Brn2+ DCN neurons (B, in red) and Tbr2+ unipolar brush cells (C, in red) in newborn control and *Olig3*$^{-/-}$ mutant mice. All cerebellar sagittal sections were counterstained with DAPI (blue).

The mean and SD are plotted in all graphs, and the dots represent the mean of individual animals. Statistical significance was determined with a two-tailed t-test, see Table S2. Photomicrographs were acquired using the automatic tile scan modus (10% overlap between tiles) of the Zeiss LSM700 confocal microscope.

**Supplementary Figure 4. Cerebellar neuron loss in *Olig3* mutant embryos.**
(**A-C**) Immunofluorescence characterization and quantification of Tbr1+ DCN neurons (A, in red), Pax6+ EGL cells (B, in red) and Foxp2+ Purkinje cells (C, in red) in E13.5 control (*Olig3*$^{GFP/+}$) and *Olig3* (*Olig3*$^{GFP/GFP}$) mutant embryos. The nuclear transitory zone (NTZ) is marked in A. All cerebellar sagittal sections were stained against GFP (green). The boxed areas on the micrographs are illustrated to the right of the main photographs with or without GFP staining for a better visualization of the Tbr1+, Pax6+ and Foxp2+ cells.
The mean and SD are plotted in all graphs, and the dots represent the mean of individual animals. Statistical significance was determined with a two-tailed t-test, see Table S2. Photomicrographs were acquired using the automatic tile scan modus (10% overlap between tiles) of the Zeiss LSM700 confocal microscope.

**Supplementary Figure 5. Deficits of rhombic progenitor cells in *Olig3* mutant embryos.**
(**A, B**) Left, immunofluorescence characterization of Atoh1+ rhombic lip cells (A, in red) and Ptf1a+ ventricular zone cells (B, in red) in control (*Olig3*$^{GFP/+}$) and *Olig3* mutant (*Olig3*$^{GFP/GFP}$) embryos at E12.5. Right, quantification of Atoh1+ (A) and Ptf1a+ (B) cells in control and Olig3 mutant mice at E11.5 and E12.5. All cerebellar sagittal sections were stained against GFP (green).
(**C**) Left, immunofluorescence characterization of proliferative BrdU+ (red) in the rhombic lip and ventricular zone of control (*Olig3*$^{GFP/+}$) and *Olig3* (*Olig3*$^{GFP/GFP}$) mutant mice at E12.5. The numbered boxed areas are displayed to the right of the main photographs. Right, quantification of BrdU+ cells in the rhombic lip (RL, upper) and ventricular zone (VZ, lower) in control and *Olig3* mutant mice at E11.5 and E12.5. All cerebellar sagittal sections were stained against GFP (green).
(**D**) Left, immunofluorescence characterization of Tunel+ apoptotic bodies (puncta, red) in control (*Olig3*$^{GFP/+}$) and *Olig3* mutant (*Olig3*$^{GFP/GFP}$) mice at E12.5. Right, quantification of Tunel+ apoptotic bodies in control and *Olig3*

mutant mice at E11.5 and E12.5. All cerebellar sagittal sections were stained against GFP (green).

The mean and SD are plotted in all graphs, and the dots represent the mean of individual animals. Significance was obtained using a one-way ANOVA followed by *post hoc* Tukey's test, see Table S2. Photomicrographs were acquired using the automatic tile scan modus (10% overlap between tiles) of the Zeiss spinning disk confocal microscope (in A and D) and the Zeiss LSM700 confocal microscope (in B and C).

**Supplementary Figure 6. Misspecified Foxp2+/Pax2+ cell numbers decline over the time in *Olig3* mutant mice.**

(**A**) Analysis of Foxp2+ (blue) and Pax2+ (red) cells in control (*Olig3*$^{GFP/+}$) and *Olig3* mutant (*Olig3*$^{GFP/GFP}$) mice at E14.5 (upper panels) and P0 (lower panels). The numbered boxed areas are displayed as magnifications to the right of the main photographs. GFP (green) expressed from the *Olig3* locus can be seen at E14.5 but no longer by P0.

(**B**) Quantification of the proportion of Pax2+ cells co-expressing Foxp2 in control and *Olig3*$^{-/-}$ mutant mice at the indicated stages.

(**C**) Analysis of E15.5 wildtype mice that were electroporated at E14.0 with control (*pCAG-GFP* + *Empty-IRES-GFP*) or Olig3-overexpression (*pCAG-Olig3-IRES-GFP*) plasmids. Representative analyzed cells imaged from the cerebellum of electroporated embryos that were stained against Pax2 (red) and GFP (green). Quantification of the proportion of GFP+ cells co-expressing Pax2 in electroporated control (Olig3-) and Olig3-overexpressing (Olig3+) mice is displayed in Figure 5C.

The mean and SD are plotted in the graph, and the dots represent the mean of individual animals. Significance was obtained using a one-way ANOVA followed by *post hoc* Tukey's test, see Table S2. Photomicrographs in A were acquired using the automatic tile scan modus (10% overlap between tiles) of the Zeiss spinning disk confocal microscope (upper panels) and the Zeiss LSM700 confocal microscope (lower panels). Photomicrographs in C were manually acquired using a Leica SPL confocal microscope.

**Supplementary Figure 7. Analysis of GABAergic neurons in O*lig2* mutant mice.**

(**A**) Immunofluorescence characterization and quantification of Foxp2+ (green) cells in control (*Olig2*$^{+/-}$) and *Olig2* mutant (*Olig2*$^{-/-}$) mice at E18.5. All cerebellar sagittal sections were counterstained with DAPI (blue).

(**B**) Immunofluorescence characterization and quantification of Pax2+ (green) cells in control and *Olig2*$^{-/-}$ mutant mice at E18.5. All cerebellar sagittal sections were counterstained with DAPI (blue).

(**C**) Analysis of cerebellar GABAergic neurons with a history of *Olig2* (βgal+) expression. *Olig2*$^{cre/+}$*;Tau*$^{nLacZ/+}$ mice were analyzed at E18.5. Sagittal sections from these mice were stained against βgal (red) and markers for Purkinje cells (Foxp2, blue in upper panel) and inhibitory interneurons (Pax2, blue in lower panel). Double-positive (βgal+/marker+) cells were quantified at E18.5.

(**D**) Summarized comparison of the percentage of GABAergic neurons with a history of *Olig3* and *Olig2* expression. See Figure 2G and H (for Olig3 analysis) and panel C above (for Olig2 analysis).

(**E**) Immunofluorescence characterization of Olig2+ (red) cells in control (*Olig3*$^{+/}$) and *Olig3* mutant (*Olig3*$^{-/-}$) mice at E12.5. For quantifications see Figure 6C.

(**F**) Immunofluorescence characterization of Olig3+ (red) cells in control (*Olig2*$^{+/-}$) and *Olig2* mutant (*Olig2*$^{-/-}$) mice at E12.5. For quantifications see Figure 6D.

The mean and SD are plotted in all graphs, and the dots represent the mean of individual animals. Statistical significance was obtained with a two-tailed t-test, see Table S2. Photomicrographs were acquired using the automatic tile scan modus (10% overlap between tiles) of the Zeiss spinning disk confocal microscope (in A, B, E and F) and the Zeiss LSM700 confocal microscope (in C).

**Video S1.**

Three-dimensional reconstruction of an E19 *Olig3*$^{creERT2/+}$*; Ai14*$^{+/-}$ mouse brain that was recombined at E10.5. Red fluorescence represents the somas and axons of all cells with a history of Olig3 expression. See also Figure S2A.

**Supplementary Table 1. Categorization of bHLH transcription factors expressed or not expressed during cerebellar development in mice.**

| **bHLH factors expressed in cerebellar progenitor niches: Rhombic lip (RL), Ventricular zone (VZ) and/or external granular cell layer (EGL)** | | | | | |
|---|---|---|---|---|---|
| Gene Name | Expressed in progenitors? | Developmental stage: embryonic (E) day | | | |
| | | E11.5 | E13.5 | E15.5 | E17.5/E18.5 |
| Ascl1 | Yes | VZ | VZ | Weak in VZ | Not expressed |
| Atoh1 | Yes | RL | RL & EGL | RL & EGL | RL & EGL |
| Hes1 | Yes | Not expressed | RL & VZ | Not expressed | Not expressed |
| Hes5 | Yes | RL & VZ | RL & VZ | RL & VZ | Postmitotic cells |
| Hes6 | Yes | RL & VZ | RL, VZ & EGL | EGL | No data |
| Hes7 | Yes | RL & VZ | Weak in RL, VZ | Weak in RL, VZ | Not expressed |
| Hey1 | Yes | Not expressed | Not expressed | EGL | EGL |
| Hif1a | Yes | RL & VZ | Weak in RL, VZ | Not expressed | Not expressed |
| Id1 | Yes | RL & VZ | In blood vessels | In blood vessels | In blood vessels |
| Id3 | Yes | RL & VZ | RL, VZ & EGL | RL, VZ & EGL | EGL |
| Max | Yes | Weak in RL & VZ | Not expressed | Not expressed | Not expressed |
| Mxd3 | Yes | RL & VZ | RL, VZ & EGL | EGL | EGL |
| Mxi1 | Yes | Weak in RL & VZ | RL, VZ & EGL | RL, VZ & EGL | Weak in EGL |
| Mycn | Yes | Strong in RL & VZ | Not expressed | Not expressed | Not expressed |
| Neurod1 | Yes | Not expressed | Not expressed | Strong in EGL | Strong in EGL |
| Neurod6 | Yes | Not expressed | Weak in VZ | VZ | Broad expression |
| Neurog1 | Yes | Not expressed | VZ | Not expressed | Not expressed |
| Neurog2 | Yes | Not expressed | Weak in VZ | Strong in VZ | Not expressed |
| Olig2 | Yes | Weak in VZ | Strong in VZ | Postmitotic cells | Postmitotic cells |
| Olig3 | Yes | RL & weak in VZ | RL & VZ | Not expressed | Not expressed |
| Ptf1a | Yes | Strong in VZ | Strong in VZ | Weak in VZ | Not expressed |
| Srebf1 | Yes | Not expressed | RL, VZ & EGL | Weak in EGL | Not expressed |
| Srebf2 | Yes | Weak in RL & VZ | RL & VZ | Postmitotic cells | Postmitotic cells |
| Tcf12 | Yes | RL & VZ | RL, VZ & EGL | RL, VZ & EGL | RL, VZ & EGL |
| Tcf3 | Yes | RL & VZ | RL, VZ & EGL | RL, VZ & EGL | RL, VZ & EGL |
| Tcf4 | Yes | RL & VZ | RL, VZ & EGL | RL, VZ & EGL | RL, VZ & EGL |

| **bHLH factors expressed in postmitotic cerebellar cells during development** | |
|---|---|
| Gene name: | Arnt2, Bhlhe22, Clock, Epas1, Id2, id4, Mlx, Mnt, Mxd1, Mxd4, Myc, Ncoa1, Ncoa2, Neurod2, Neurog3, Nhlh1, Nhlh2, Npas3, Npas4, Olig1, Scx, Sim2, Usf1 & Usf2 |

| **bHLH factors not expressed in the cerebellum during development** | |
|---|---|
| Gene name: | Ahr, Ahrr, Arnt, Arntl2, Ascl2, Ascl3, Ascl4, Ascl5, Atoh7, Atoh8, Bhlha15, Bhlha9, Bhlhb9, Bhlhe23, Bhlhe40, Bhlhe41, Ferd3l, Figla, Hand1, Hand2, Helt, Hes2, Hes3, Hes4, Hey2, Heyl, Hif3a, Lyl1, Mesp1, Mesp2, Mitf, Mlxip, Mlxipl, Msc, Myf5, Myf6, Myod1, Myog, Ncoa3, Neurod4, Npas1, Npas2, Sim1, Sohlh1, Sohlh2, Tal1, Tal2, Tcf15, Tcf21, Tcf23, Tcf24, Tcfl5, Tfap4, Tfe3, Tfeb, Tfec, Twist1 & Twist2 |

#### Supplementary Table 2. Description of the statistical analyses used in this study.

| Fig. | n | Descriptive statistics | Test used | P value | Degrees of freedom and F/t/z/R/ETC | Pos hoc analysis | Adjusted p value |
|---|---|---|---|---|---|---|---|
| 2E | 3 mice (E10.5)<br>3 mice (E11.5)<br>3 mice (E12.5)<br>3 mice (E13.5) | Mean and SD | Ordinary one-way ANOVA | <0.0001 | F: 67.20<br>F(DFn, DFd): 0.1441 (3, 8) | Tukey's multiple comparative test | As indicated in the figure |
| 2F | 3 mice (E10.5)<br>3 mice (E11.5)<br>3 mice (E12.5)<br>3 mice (E13.5) | Mean and SD | Ordinary one-way ANOVA | <0.0001 | F: 93.57<br>F(DFn, DFd): 1.868 (3, 8) | Tukey's multiple comparative test | As indicated in the figure |
| 2G | 3 mice (E10.5)<br>3 mice (E11.5)<br>3 mice (E12.5)<br>3 mice (E13.5) | Mean and SD | Ordinary one-way ANOVA | <0.0001 | F: 122.2<br>F(DFn, DFd): 1.096 (3, 8) | Tukey's multiple comparative test | As indicated in the figure |
| 2H | 3 mice (E10.5)<br>3 mice (E11.5)<br>3 mice (E12.5)<br>3 mice (E13.5) | Mean and SD | Ordinary one-way ANOVA | <0.0001 | F: 13.65<br>F(DFn, DFd): 0.8851 (3, 8) | Tukey's multiple comparative test | As indicated in the figure |
| 3A | 4 control mice<br>4 mutant mice | Mean and SD | Unpaired t-test (two-tailed) | <0.0001 | t=15.13; df=6 | - | - |
| 3B | 4 control mice<br>4 mutant mice | Mean and SD | Unpaired t-test (two-tailed) | 0.0005 | t=6.742; df=6 | - | - |
| 3C | 3 control mice<br>3 mutant mice | Mean and SD | Unpaired t-test (two-tailed) | 0.0005 | t=10.49; df=4 | - | - |
| 3D | 3 control mice<br>3 mutant mice | Mean and SD | Unpaired t-test (two-tailed) | 0.0002 | t=13.43; df=4 | - | - |
| 3E | 4 control mice<br>4 mutant mice | Mean and SD | Unpaired t-test (two-tailed) | <0.0001 | t=16.89; df=6 | - | - |
| 3F | 4 control mice<br>4 mutant mice | Mean and SD | Unpaired t-test (two-tailed) | <0.0001 | t=11.28; df=6 | - | - |
| 4D | 4 control (E13.5)<br>4 mutant (E13.5)<br>4 control (E14.5)<br>4 mutant (E14.5)<br>4 control (P0)<br>4 mutant (P0) | Mean and SD | Ordinary one-way ANOVA | <0.0001 | F: 499.1<br>F(DFn, DFd): 1.245 (5, 18) | Tukey's multiple comparative test | As indicated in the figure |
| 4D | 4 control (E13.5)<br>4 mutant (E13.5)<br>4 control (E14.5)<br>4 mutant (E14.5)<br>4 control (P0)<br>4 mutant (P0) | Mean and SD | Ordinary one-way ANOVA | <0.0001 | F: 514.2<br>F(DFn, DFd): 7.873 (5, 18) | Tukey's multiple comparative test | As indicated in the figure |
| 4F | 4 control mice<br>3 mutant mice | Mean and SD | Unpaired t-test (two-tailed) | <0.0001 | t=22.92; df=5 | - | - |
| 5B | 3 control mice<br>7 Pax2-OE mice | Mean and SD | Unpaired t-test (two-tailed) | <0.0001 | t=67.67; df=8 | - | - |
| 5C | 5 control mice<br>5 Olig3-OE mice | Mean and SD | Unpaired t-test (two-tailed) | 0.0001 | t=13.24; df=8 | - | - |
| 6C | 3 control mice<br>3 mutant mice | Mean and SD | Unpaired t-test (two-tailed) | 0.9102 | t=0.1201; df=4 | - | - |
| 6D | 3 control mice<br>3 mutant mice | Mean and SD | Unpaired t-test (two-tailed) | 0.6912 | t=0.4272; df=4 | - | - |
| 6D | 4 control<br>4 Olig3 mutant<br>4 Olig2 mutant | Mean and SD | Ordinary one-way ANOVA | <0.0001 | F: 1443<br>F(DFn, DFd): 0.7101 (2, 11) | Tukey's multiple comparative test | As indicated in the figure |

| | | | | | | | |
|---|---|---|---|---|---|---|---|
| S2C | 3 mice (E10.5)<br>3 mice (E11.5)<br>3 mice (E12.5)<br>3 mice (E13.5) | Mean and SD | Ordinary one-way ANOVA | <0.0001 | F: 299.8<br>F(DFn, DFd): 0.3155 (3, 8) | Tukey's multiple comparative test | As indicated in the figure |
| S2D | 3 mice (E10.5)<br>3 mice (E11.5)<br>3 mice (E12.5)<br>3 mice (E13.5) | Mean and SD | Ordinary one-way ANOVA | <0.0001 | F: 190.0<br>F(DFn, DFd): 0.8347 (3, 8) | Tukey's multiple comparative test | As indicated in the figure |
| S2E | 3 mice (E10.5)<br>3 mice (E11.5)<br>3 mice (E12.5)<br>3 mice (E13.5) | Mean and SD | Ordinary one-way ANOVA | <0.0001 | F: 85.64<br>F(DFn, DFd): 0.08333 (3, 8) | Tukey's multiple comparative test | As indicated in the figure |
| S3B | 3 control mice<br>3 mutant mice | Mean and SD | Unpaired t-test (two-tailed) | 0.0092 | t=4.714; df=4 | - | - |
| S3C | 3 control mice<br>3 mutant mice | Mean and SD | Unpaired t-test (two-tailed) | 0.2731 | t=1.270; df=4 | - | - |
| S4A | 3 control mice<br>3 mutant mice | Mean and SD | Unpaired t-test (two-tailed) | <0.0001 | t=17.21; df=4 | - | - |
| S4B | 3 control mice<br>3 mutant mice | Mean and SD | Unpaired t-test (two-tailed) | <0.0001 | t=17.07; df=4 | - | - |
| S4C | 3 control mice<br>3 mutant mice | Mean and SD | Unpaired t-test (two-tailed) | 0.0803 | t=2.330; df=4 | - | - |
| S5A | 4 control (E11.5)<br>4 mutant (E11.5)<br>4 control (E12.5)<br>4 mutant (E12.5) | Mean and SD | Ordinary one-way ANOVA | <0.0001 | F: 141.0<br>F(DFn, DFd): 2.264 (3, 12) | Tukey's multiple comparative test | As indicated in the figure |
| S5B | 3 control (E11.5)<br>4 mutant (E11.5)<br>4 control (E12.5)<br>4 mutant (E12.5) | Mean and SD | Ordinary one-way ANOVA | <0.0001 | F: 119.2<br>F(DFn, DFd): 0.3046 (3, 11) | Tukey's multiple comparative test | As indicated in the figure |
| S5C (RL) | 3 control (E11.5)<br>4 mutant (E11.5)<br>3 control (E12.5)<br>4 mutant (E12.5) | Mean and SD | Ordinary one-way ANOVA | <0.0001 | F: 228.7<br>F(DFn, DFd): 0.4398 (3, 10) | Tukey's multiple comparative test | As indicated in the figure |
| S5C (VZ) | 4 control (E11.5)<br>4 mutant (E11.5)<br>4 control (E12.5)<br>4 mutant (E12.5) | Mean and SD | Ordinary one-way ANOVA | <0.0001 | F: 156.8<br>F(DFn, DFd): 0.9683 (3, 12) | Tukey's multiple comparative test | As indicated in the figure |
| S5D | 4 control (E11.5)<br>4 mutant (E11.5)<br>4 control (E12.5)<br>4 mutant (E12.5) | Mean and SD | Ordinary one-way ANOVA | 0.1739 | F: 1.960<br>F(DFn, DFd): 0.4100 (3, 12) | Tukey's multiple comparative test | As indicated in the figure |
| S6B | 4 control (E13.5)<br>4 mutant (E13.5)<br>4 control (E14.5)<br>4 mutant (E14.5)<br>4 control (P0)<br>4 mutant (P0) | Mean and SD | Ordinary one-way ANOVA | <0.0001 | F: 850.2<br>F(DFn, DFd): 3.630 (5, 18) | Tukey's multiple comparative test | As indicated in the figure |
| S7A | 3 control mice<br>4 mutant mice | Mean and SD | Unpaired t-test (two-tailed) | <0.0001 | t=12.83; df=5 | - | - |
| S7B | 3 control mice<br>4 mutant mice | Mean and SD | Unpaired t-test (two-tailed) | 0.0002 | t=10.00; df=5 | - | - |

Abbreviation: OE, overexpression.

Figure S1

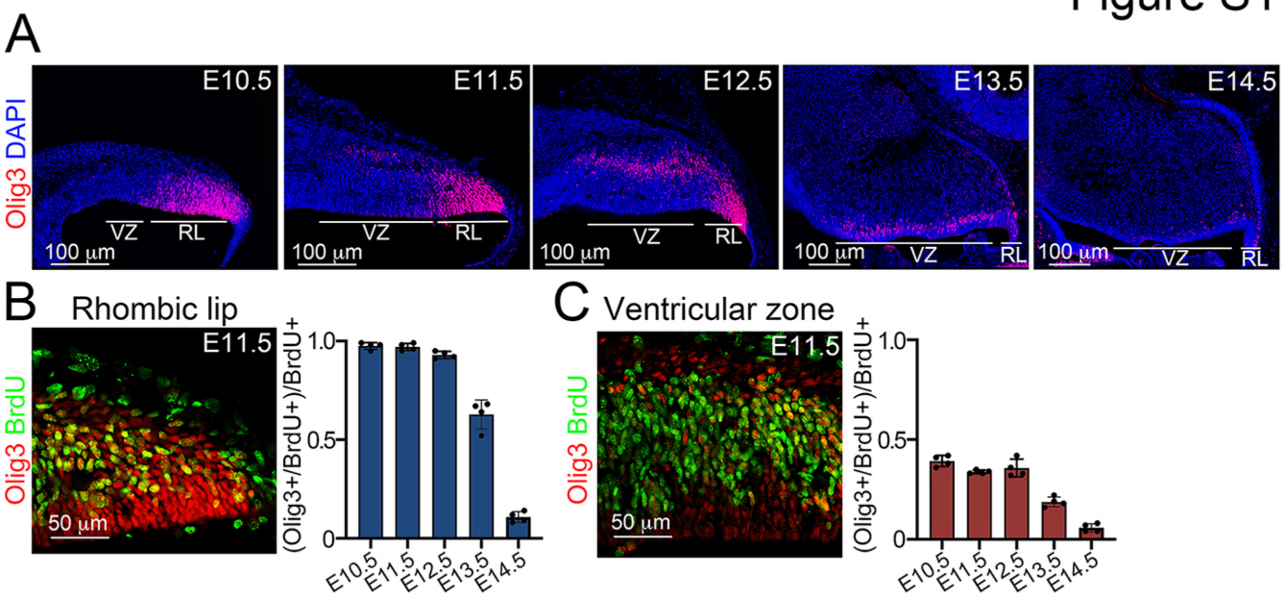
A
Olig3 DAPI
E10.5
E11.5
E12.5
E13.5
E14.5
VZ
RL
100 μm
B
Rhombic lip
Olig3 BrdU
E11.5
50 μm
(Olig3+/BrdU+)/BrdU+
1.0
0.5
0
E10.5
E11.5
E12.5
E13.5
E14.5
C
Ventricular zone

### Figure S2

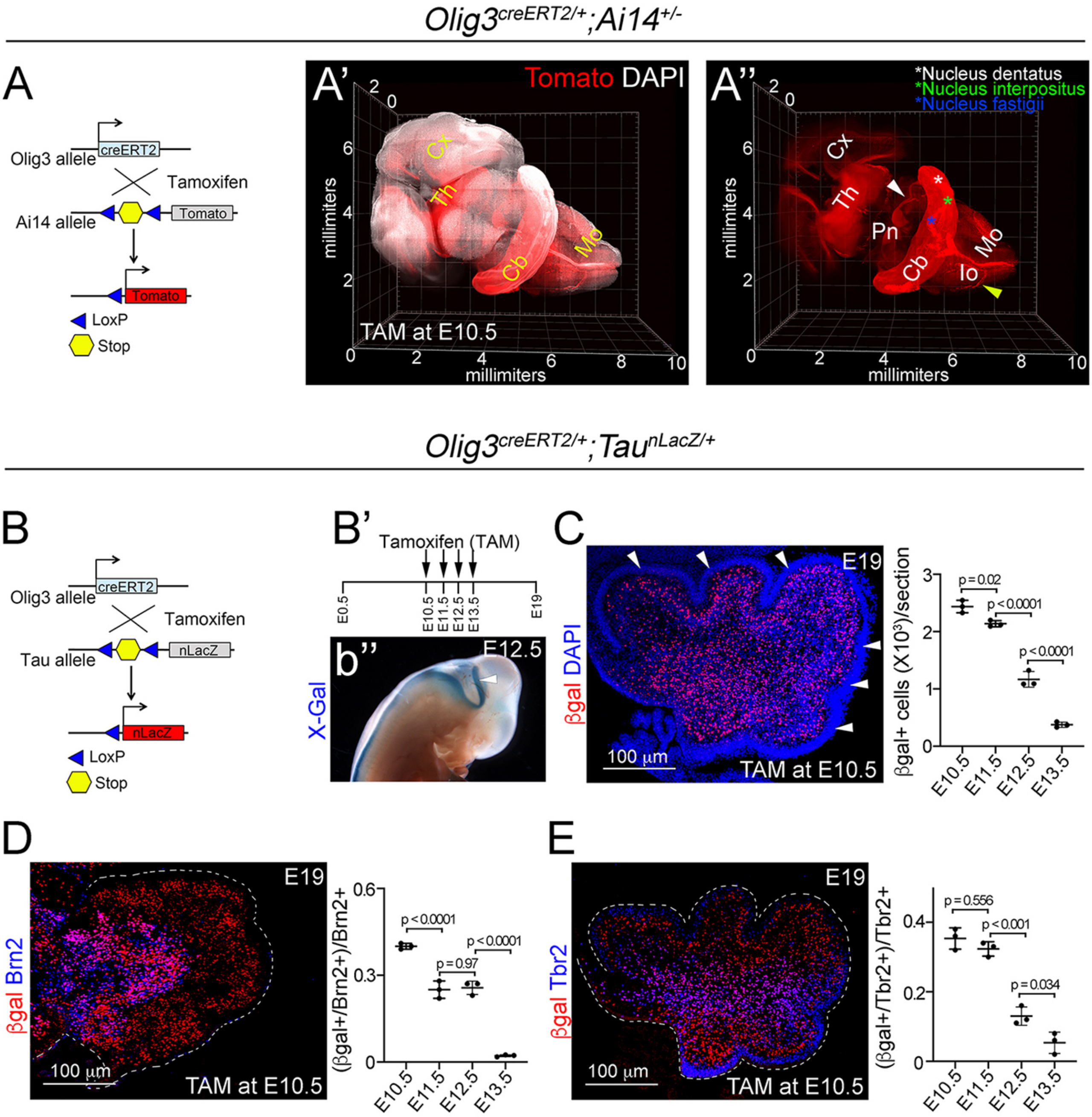
Olig3creERT2/+;Ai14+/-
A
Olig3 allele
creERT2
Tamoxifen
Ai14 allele
Tomato
Tomato
LoxP
Stop
A'
Tomato DAPI
Cx
Th
Cb
Mo
millimiters
TAM at E10.5
A''
*Nucleus dentatus
*Nucleus interpositus
*Nucleus fastigii
Cx
Th
Pn
Cb
Mo
Io
millimiters
Olig3creERT2/+;TaunLacZ/+
B
Olig3 allele
creERT2
Tamoxifen
Tau allele
nLacZ
nLacZ
LoxP
Stop
B'
Tamoxifen (TAM)
E0.5
E10.5
E11.5
E12.5
E13.5
E19
b''
E12.5
X-Gal
C
E19
βgal DAPI
100 μm
TAM at E10.5
βgal+ cells (X10³)/section
p = 0.02
p < 0.0001
p < 0.0001
E10.5
E11.5
E12.5
E13.5
D
E19
βgal Brn2
100 μm
TAM at E10.5
(βgal+/Brn2+)/Brn2+
p < 0.0001
p = 0.97
p < 0.0001
E10.5
E11.5
E12.5
E13.5
E
E19
βgal Tbr2
100 μm
TAM at E10.5
(βgal+/Tbr2+)/Tbr2+
p = 0.556
p < 0.001
p = 0.034
E10.5
E11.5
E12.5
E13.5

### Figure S3

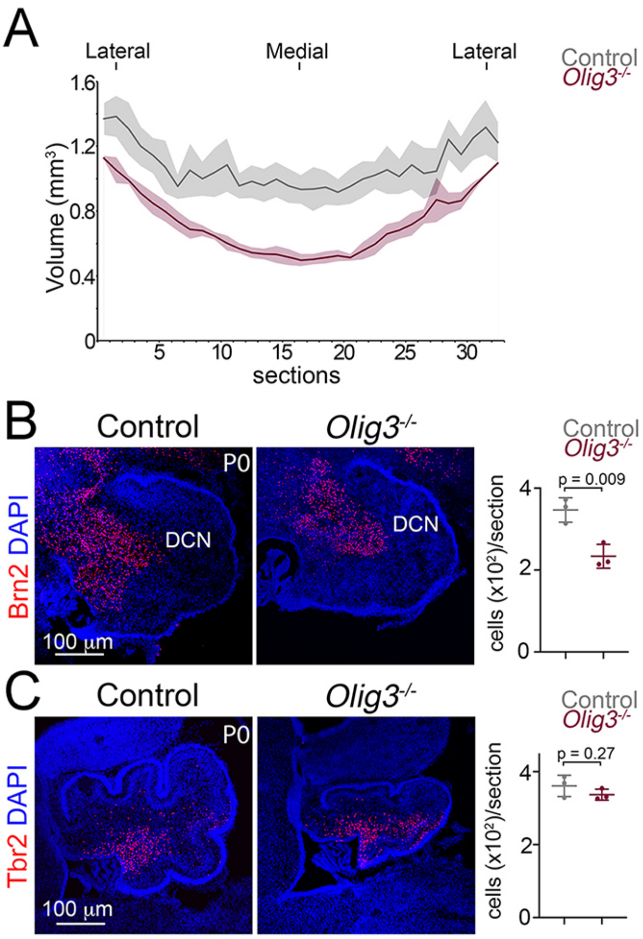
A
Lateral
Medial
Lateral
Control
Olig3-/-
Volume (mm3)
1.6
1.2
0.8
0.4
0
5
10
15
20
25
30
sections
B
Control
Olig3-/-
P0
Brn2 DAPI
DCN
DCN
100 μm
Control
Olig3-/-
cells (x10²)/section
p = 0.009
4
2
0
C
Control
Olig3-/-
P0
Tbr2 DAPI
100 μm
Control
Olig3-/-
cells (x10²)/section
p = 0.27
4
2
0

### Figure S4

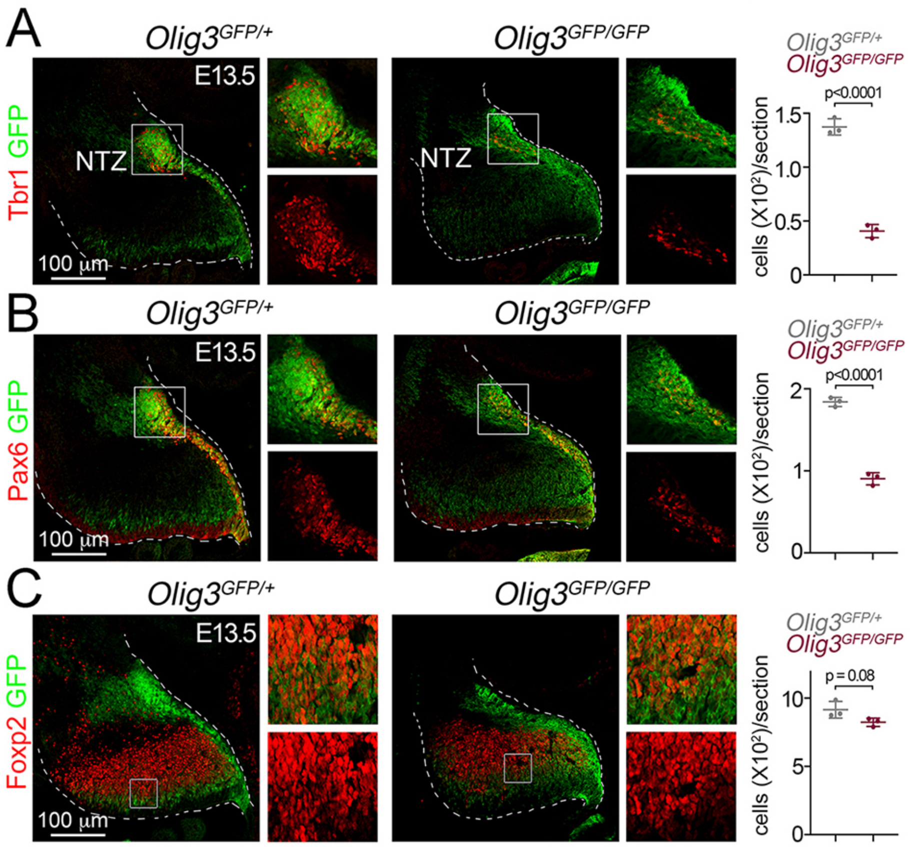

A
Olig3GFP/+
Olig3GFP/GFP
Tbr1 GFP
E13.5
NTZ
NTZ
100 µm
Olig3GFP/+
Olig3GFP/GFP
p<0.0001
cells (X10²)/section
1.5
1.0
0.5
0
B
Olig3GFP/+
Olig3GFP/GFP
Pax6 GFP
E13.5
100 µm
Olig3GFP/+
Olig3GFP/GFP
p<0.0001
cells (X10²)/section
2
1
0
C
Olig3GFP/+
Olig3GFP/GFP
Foxp2 GFP
E13.5
100 µm
Olig3GFP/+
Olig3GFP/GFP
p = 0.08
cells (X10²)/section
10
5
0

### Figure S5

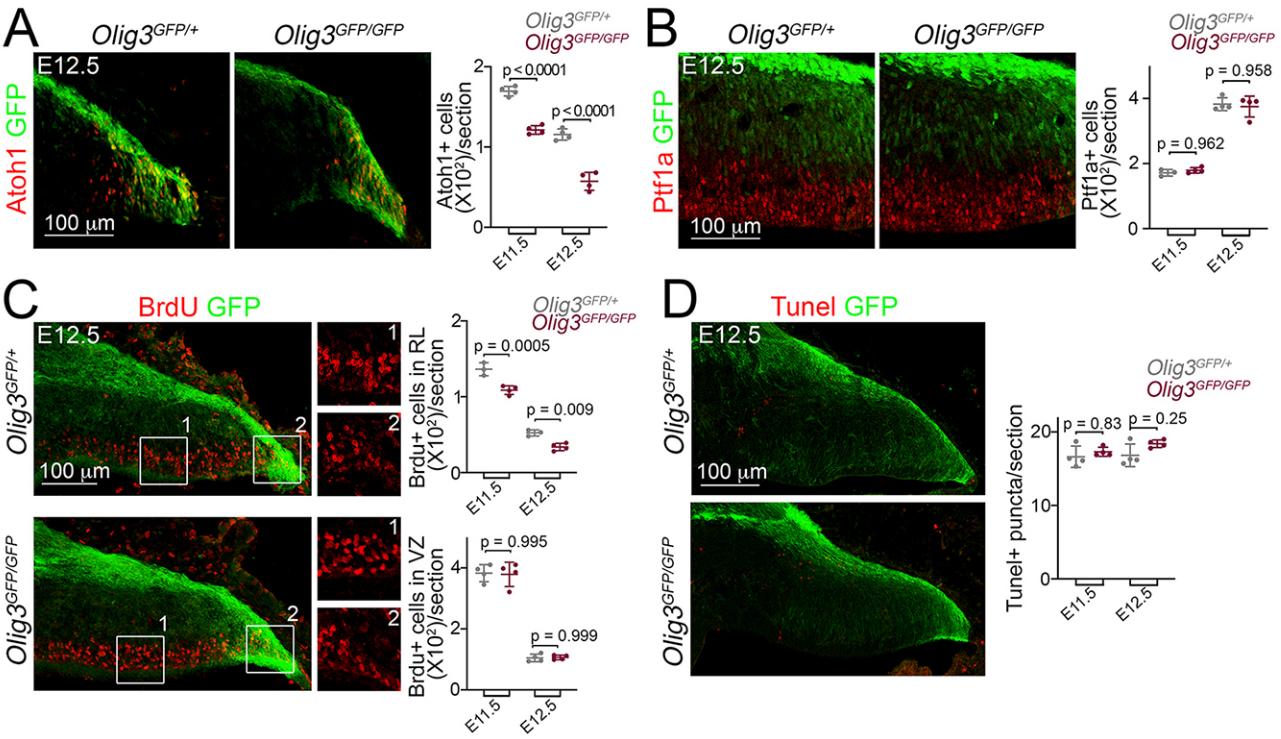

A
Olig3GFP/+
Olig3GFP/GFP
E12.5
Atoh1 GFP
100 µm
Olig3GFP/+
Olig3GFP/GFP
p < 0.0001
p < 0.0001
Atoh1+ cells (X10²)/section
E11.5
E12.5
B
Olig3GFP/+
Olig3GFP/GFP
E12.5
Ptf1a GFP
100 µm
p = 0.958
p = 0.962
Ptf1a+ cells (X10²)/section
E11.5
E12.5
C
BrdU GFP
E12.5
Olig3GFP/+
Olig3GFP/GFP
100 µm
p = 0.0005
p = 0.009
Brdu+ cells in RL (X10²)/section
p = 0.995
p = 0.999
Brdu+ cells in VZ (X10²)/section
E11.5
E12.5
D
Tunel GFP
E12.5
Olig3GFP/+
Olig3GFP/GFP
100 µm
p = 0.83
p = 0.25
Tunel+ puncta/section
E11.5
E12.5

Figure S6

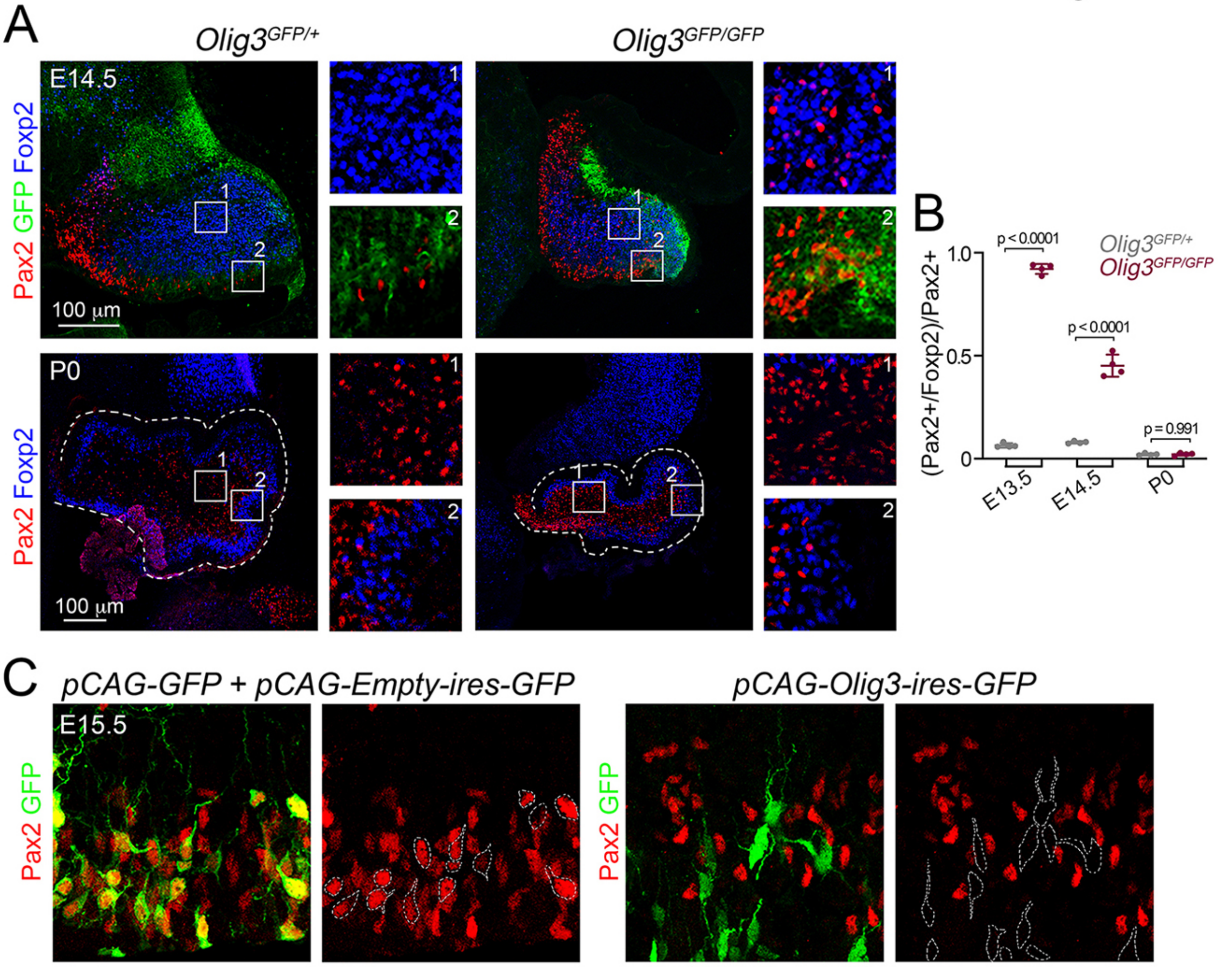
A
Olig3GFP/+
Olig3GFP/GFP
E14.5
Pax2 GFP Foxp2
100 μm
P0
Pax2 Foxp2
100 μm
B
(Pax2+/Foxp2)/Pax2+
1.0
0.5
0
p < 0.0001
p < 0.0001
p = 0.991
Olig3GFP/+
Olig3GFP/GFP
E13.5
E14.5
P0
C
pCAG-GFP + pCAG-Empty-ires-GFP
pCAG-Olig3-ires-GFP
E15.5
Pax2 GFP

### Figure S7

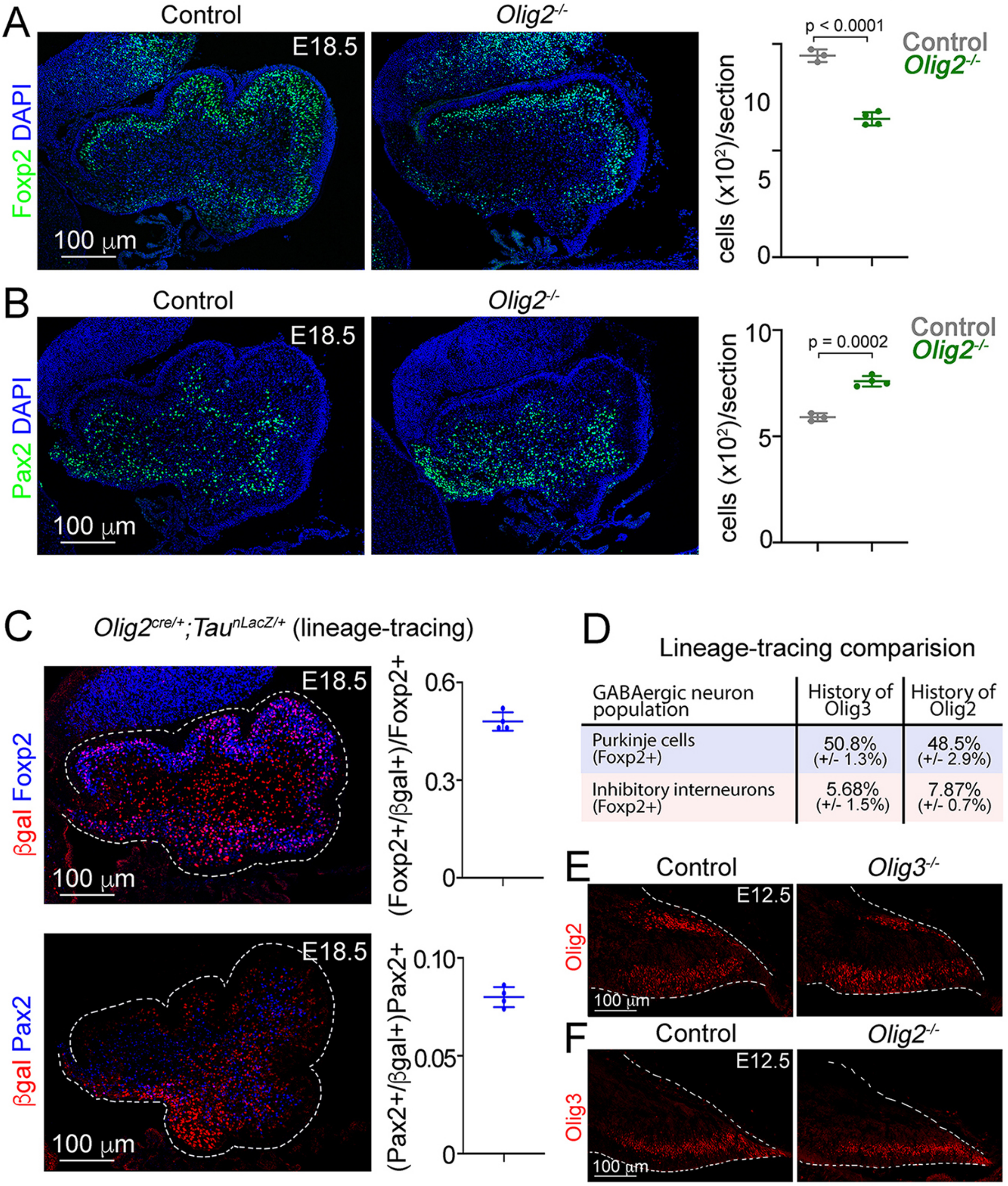
A
Control
Olig2-/-
E18.5
Foxp2 DAPI
100 μm
p < 0.0001
Control
Olig2-/-
cells (x10²)/section
0
5
10
B
Control
Olig2-/-
E18.5
Pax2 DAPI
100 μm
p = 0.0002
Control
Olig2-/-
cells (x10²)/section
0
5
10
C
Olig2cre/+;TaunLacZ/+ (lineage-tracing)
E18.5
βgal Foxp2
100 μm
(Foxp2+/βgal+)/Foxp2+
0
0.3
0.6
E18.5
βgal Pax2
100 μm
(Pax2+/βgal+)Pax2+
0
0.05
0.10
D
Lineage-tracing comparision
GABAergic neuron population | History of Olig3 | History of Olig2
Purkinje cells (Foxp2+) | 50.8% (+/- 1.3%) | 48.5% (+/- 2.9%)
Inhibitory interneurons (Foxp2+) | 5.68% (+/- 1.5%) | 7.87% (+/- 0.7%)
E
Control
Olig3-/-
E12.5
Olig2
100 μm
F
Control
Olig2-/-
E12.5
Olig3
100 μm
